# A novel human model to deconvolve cell-intrinsic phenotypes of genetically dysregulated pathways in lung squamous cell carcinoma

**DOI:** 10.1101/2023.12.13.568969

**Authors:** Julia Ogden, Robert Sellers, Sudhakar Sahoo, Anthony Oojageer, Anshuman Chaturvedi, Caroline Dive, Carlos Lopez-Garcia

**Affiliations:** Cancer Research UK Manchester Institute, Wilmslow Road, M20 4BX, Manchester (United Kingdom); Department of Histopathology, The Christie Hospital, Wilmslow Road, Manchester, M20 4BX (United Kingdom); Cancer Research UK, National Biomarker Centre, Wilmslow Road, M20 4BX, Manchester (United Kingdom); Cancer Research UK Lung Cancer Centre of Excellence, Wilmslow Road, M20 4BX, Manchester (United Kingdom)

## Abstract

Tractable, patient relevant models are needed to investigate cancer progression and heterogeneity. Here, we report an alternative and unique in vitro model of lung squamous cell carcinoma (LUSC) using primary human bronchial epithelial cells (hBECs) from three healthy donors. The co-operation of ubiquitous alterations (*TP53* and *CDKN2A* loss) and components of commonly deregulated pathways including squamous differentiation (*SOX2*), PI3K signalling (*PTEN*) and the oxidative stress response (*KEAP1*) was investigated by generating hBECs harbouring cumulative alterations. Our analyses confirmed that *SOX2*-overexpression initiates early preinvasive LUSC stages, and co-operation with the oxidative stress response and PI3K pathways to drive more aggressive phenotypes, with expansion of cells expressing LUSC biomarkers and invasive properties. This cooperation was consistent with the classical LUSC subtype. Importantly, we connected pathway dysregulation with gene expression changes associated with cell-intrinsic processes and immunomodulation. Our approach constitutes a powerful system to model LUSC and unravel genotype-phenotype causations of clinical relevance.

## BACKGROUND

Lung cancers are a devastating group of neoplasms which broadly divide into two categories, small cell lung cancer (SCLC) and non-small cell lung cancer (NSCLC). NSCLC accounts for 85% of lung cancer cases and primarily consists of two histologically distinct subtypes including lung adenocarcinoma (LUAD) (50% of all lung cancers) and lung squamous cell carcinoma (LUSC) (30% of all lung cancers) (1). LUSC is a complex disease which predominantly originates in the proximal airways. Basal cells, which function as the multipotent progenitors of the bronchial epithelium, are believed to be the most likely cells-of-origin (2), although a more complex interplay between genetic drivers and cell-of-origin has been observed (3, 4).

Contrasting with LUAD, patients with LUSC are almost exclusively smokers, their survival is poorer and targeted treatments have largely failed (5–7). LUSC exhibits a heterogeneous landscape of phenotypic subtypes with unclear biological origins (4, 8). Genetically, LUSC is characterised by inactivation of *TP53* and upregulation of the CDK4/6 pathway (typically by *CDKN2A* inactivation) in most cases (8, 9), and unlike LUAD, dysregulation of oncogenic pathways in LUSC is less clear (9). LUSC genetic landscapes have revealed components of the squamous differentiation (SD) pathway, PI3K/Akt signalling, and oxidative stress response (OSR) pathway to be the most frequently targeted, although co-occurrence or mutual exclusivity patterns in the dysregulation of these common LUSC pathways were not observed (9). More recent multi-platform characterisations of LUSC have shown characteristic genomic features in specific subtypes (8, 10). Modelling this complexity is required to understand the underpinning biology, the specific vulnerabilities associated with each subgroup and to identify essential components of LUSC development.

Modelling LUSC heterogeneity *in vivo* has the obvious advantage of encompassing the tumour microenvironment (TME), but it is arduous, time-consuming, and costly. Non-immortalised primary human bronchial epithelial cells (hBECs), are a suitable and infrequently explored alternative. They can be expanded *in vitro* as basal cells and recapitulate bronchial epithelial architecture using organotypic systems (11). Regardless of the absence of an autochthonous TME, hBECs are an alternative to mouse models in the deconvolution of complex genotypes to find genotype-phenotype causation, identification of the essential alterations driving the cell-autonomous biology of LUSC development, uncover epistatic interactions between somatic events, and downstream actionable pathways with therapeutic application. In this report, we aimed to establish the potential of genetically engineered hBECs from three healthy donors to model LUSC and deconvolve the cell-intrinsic mechanisms whereby somatic events fuel LUSC progression using hBECs from three donors. We confirmed that the SD, PI3K/Akt and OSR pathways were all necessary for LUSC development and invasive progression, identified pro- and anti-oncogenic interactions between those pathways and matched somatic alterations with dysfunction of biologically and therapeutically actionable pathways.

### hBEC genetic-engineering strategy to model LUSC

To determine the functions and cooperation of the most frequently dysregulated pathways (Fig. 1a) in LUSC development, we designed a combinatorial strategy in which a regulator of each pathway (*SOX2*, *PTEN* and *KEAP1*) was targeted individually, or in combinations of two and three (Fig. 1a). As *TP53* mutations and CDK4/6 pathway activation (typically by *CDKN2A* loss) (8, 9) are ubiquitous in LUSC, we included concomitant *TP53*^-^and *CDKN2A* (TC) truncations (Fig. 1a). To capture some inter-person variation and avoid smoking-induced pre-existing alterations, mutants were generated using hBECs from three never-smoking donors without reported airways disease (Supplementary Table 1). Genetic manipulation was achieved by electroporation with multiplex CRISPR-Cas9 constructs, selecting *TP53*^-/-^ cells with Nutlin-3A, and lentiviral transduction to elevate the expression of *SOX2* and mimic its amplification (Fig. 1b, Supplementary Table 2 and Extended Data Fig. 1a and b). This strategy results in mutant hBECs with up to five modifications (Fig. 1a). CRISPR efficiency analysis revealed 96-100% indel efficiency and almost complete abrogation of the protein (Fig. 1c-f). Lentiviral *SOX2* resulted in *SOX2* overexpression (Fig. 1g) and analysis of SD, PI3K/Akt and OSR pathway surrogates demonstrated the expected pathway activation (Fig. 1e, f, h). We tested nine *PTEN* sgRNAs (Supplementary Table 2), only one successfully targeted the locus and this sgRNA also targets *PTENP1* (Extended Data Fig. 1c), a pseudogene with 98% *PTEN* homology. *PTENP1* encodes a lncRNA with miRNA decoy functions (12). However, as *PTENP1* indels are restricted to *PTEN* mutants and do not disrupt miRNA binding sequences, we did not anticipate any functional consequences. Analysis of the top-5 predicted off-target sites for the remaining other sgRNAs did not show any indels.

**Figure 1.**
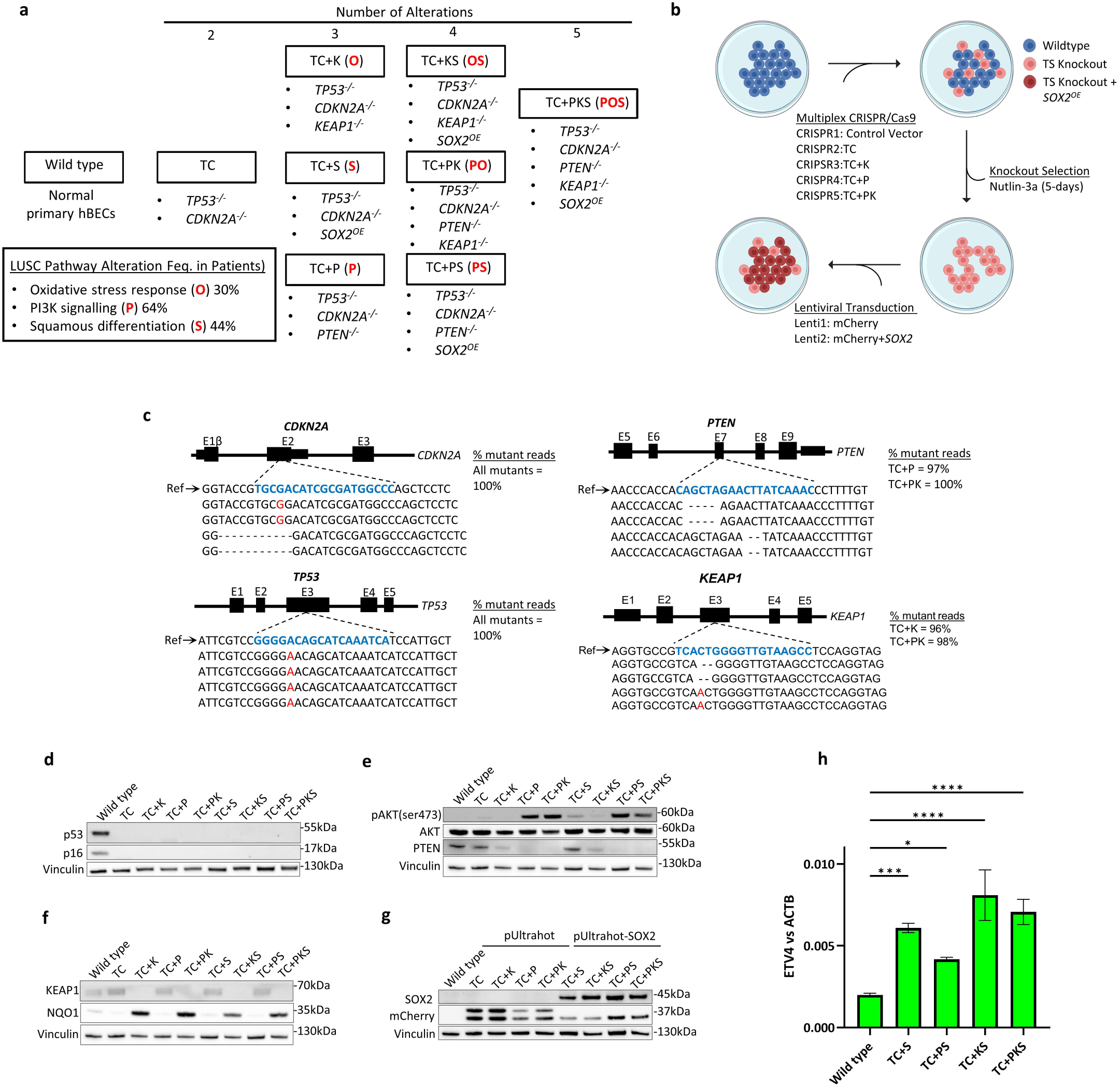
Experimental design and genetic modification strategy of hBECs. **a**. The combinatorial approach to modelling LUSC pathway dysregulation in primary hBECs. Eight mutant groups with increasing numbers of alterations were generated. All mutants harboured disruption of the *TP53* and *CDKN2A* genes. *KEAP1*, *PTEN* and *SOX2* alterations were included to target the oxidative stress response, PI3K signalling and squamous differentiation pathways, respectively. T= *TP53*, C=*CDKN2A*, P=*PTEN*, K=*KEAP1*, S=*SOX2*. Red letters indicate the pathways targeted in each mutant. O = the oxidative stress response, P = PI3K signalling, and S = squamous differentiation. **b.** Gene editing strategy for the disruption of tumour suppressor genes by electroporation with multiplex CRISPR/Cas9 vectors and lentiviral mediated *SOX2* overexpression. Five electroporations were carried out per donor: Control vector = CRISPR/cas9 vector lacking gRNA, TC=*TP53/CDKN2A* gRNAs, TC+P=*TP53/CDKN2A/PTEN* gRNAs, TC+K=*TP53/CDKN2A/KEAP1* gRNAs, TC+PK=*TP53/CDKN2A/PTEN/KEAP1* gRNAs. Those mutants with *SOX2*^OE^ were transduced with the pUltrahot vector carrying an mCherry reporter and *SOX2* cDNA and those without *SOX2*^OE^ were transduced with the empty pUltrahot vector with mCherry reporter. **c.** Next generation amplicon sequencing of CRISPR/Cas9 target loci for each gene of interest. Example mutant reads are aligned to a wild type reference sequence. gRNA target sequences are identified in blue. Red text indicates nucleotide insertions. Deleted single nucleotides are indicated (-). Right panel displays % mutant reads. **d-f.** Tumour suppressor knockouts were confirmed by western blotting for target genes. pAKT(ser473) and NQO1 were used as surrogates for activation of PI3K signalling and the oxidative stress response, respectively. **g.** Western blotting confirmed mCherry and SOX2 protein expression following transduction of hBECs with the empty pUltrahot vector, or pUltrahot with SOX2 cDNA (pUltrahot-*SOX2*). h. qPCR analysis of the *SOX2* target *ETV4* in mutants with SOX2 overexpression.

### *SOX2* inhibits proliferation in 2D cultures but enhances anchorage-independent growth and invasiveness in cooperation with the OSR and PI3/Akt pathway

Firstly, we assessed the transforming potential of our different mutant combinations in the three donors by investigating changes in proliferation, anchorage-independent growth, and invasiveness. Overall, mutants without lentiviral *SOX2*^OE^ proliferated 2-3 fold faster than wild type hBECs and *PTEN* truncations enhanced this phenotype (Fig. 2a). However, *SOX2*^OE^ mutants grew with similar rates to wild type, excluding TC+PS in donors 1 and 2 (Fig. 2a). As *SOX2*^OE^ was not associated with increased apoptosis (Extended Data Fig. 1d), this result was likely due to reduced proliferation in culture. Although paradoxical, *SOX2*^OE^ mediated growth suppression has been observed in other systems (13, 14) and indicates that SOX2 drives oncogenesis by mechanisms unrelated to proliferation. Indeed, only mutants with *SOX2*^OE^ showed detectable anchorage-independent growth over background with TC+PKS displaying maximal colony formation in soft agar (Fig. 2b, c), demonstrating that SOX2^OE^ is required for anoikis override and synergises with *PTEN* and *KEAP1* truncations. However, in donor 3, colony formation in TC+KS was comparable to TC+PKS. When we carried out invasion assays using the collagen disc method (15), we found that TC+PKS mutants showed the most invasive behaviour in all the donors (Fig. 2d, e), consistent with our soft-agar results (Fig. 2b, c). These results are in line with published *in vivo* evidence that highlighted the essential role of *SOX2* in LUSC and indicate an oncogenic role independent of proliferation. Furthermore, our results confirm the cooperation with activation of the PI3K/Akt and OSR pathways. Overall, these data characterising the newly generated models reflect carcinogenic transformation of hBECs and encouraged further analyses.

**Figure 2.**
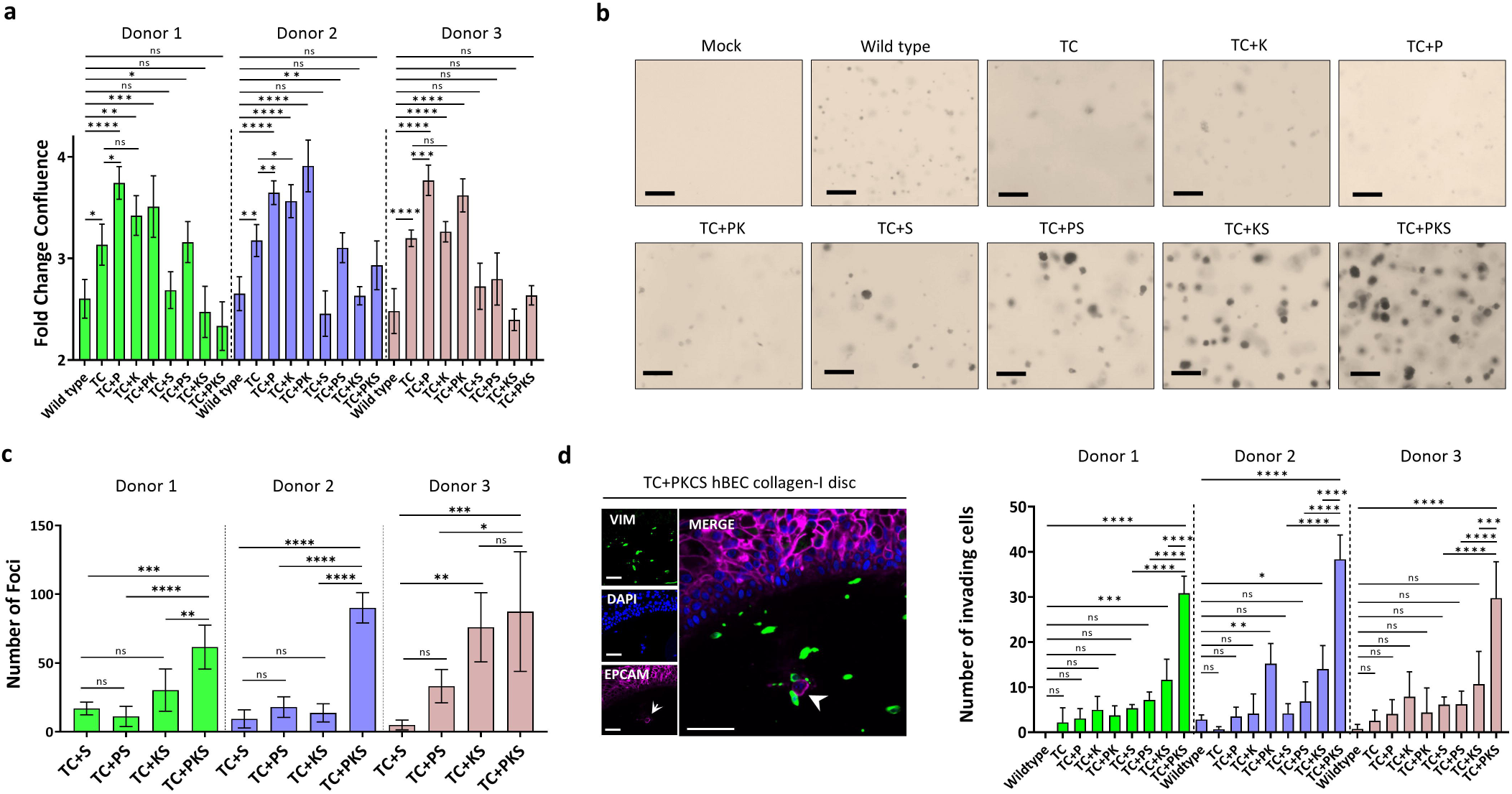
Phenotypic analysis of mutant hBECs. **a.** Cell population growth assays for wild type and mutant hBECs in three donors. Data is shown as the fold-change in confluence from 24-72 hours. Mean +/-SD (n=4). *Adj.P* values were calculated by one-way ANOVA with multiple comparisons and Holm-Šídák’s *post hoc* test. ns = not significant. **b.** Representative images of mutant hBEC colony formation in soft agar. **c.** Quantification of soft agar colonies. Foci were counted as a colony if the circumference exceeded 50µM. Mean+/-SD (n=5). Mutants with >1 colonies are shown. *Adj.P* values were calculated by one-way ANOVA with multiple comparisons and Tukey’s post *hoc test*. ns = not significant. **d.** Immunofluorescence staining of collagen-I invasion assays for the fibroblast marker vimentin and the epithelial cell marker EPCAM was used to distinguish pulmonary fibroblasts from invading epithelial cells. Left panel shows single channel images for vimentin (VIM, green), DAPI (DAPI, blue), and EPCAM (EPCAM, magenta). Right panel shows the number of EPCAM^+^ cells present in collagen. EPCAM^+^ cells were counted if ≥100µm from collagen surface. Scale bars = 50µm. Mean+/-SD of three replicates. ns = not significant.

### Effect of pathway dysregulation in bronchial epithelial homeostasis

LUSC developmental stages are characterised by epithelial changes that include loss of specialised bronchial cells, surface maturation (squamous cells) and expansion of p63-expressing cells (Fig. 3a). To ascertain the role of pathway dysregulation in the development of these epithelial perturbations, we used organotypic air-liquid interface (ALI) cultures (Figure 3b), widely used to study bronchial epithelial biology. Histological analysis revealed characteristic epithelial alterations in ALI cultures (Figure 3c-f and Extended Data Fig. 2a, b). In general, wild-type and TC mutants lacked significant differences in epithelial structure and relative abundance of specialized bronchial lineages (goblet, ciliated and club cells) (Fig. 3c-e and Extended Data Fig. 2a, b). TC+P mutants showed a prominent phenotype characterised by increased cellularity and development of intra-epithelial cysts (Figure 3c and Extended Data Fig. 2a), although without relative changes in differentiated bronchial lineages (Fig. 3 d-e). TC+K and TC+PK mutants exhibited an overall reduction in the fraction of club cells (Fig. 3e), which suggests a previously unreported function of the OSR pathway in the differentiation dynamics of that lineage.

**Figure 3.**
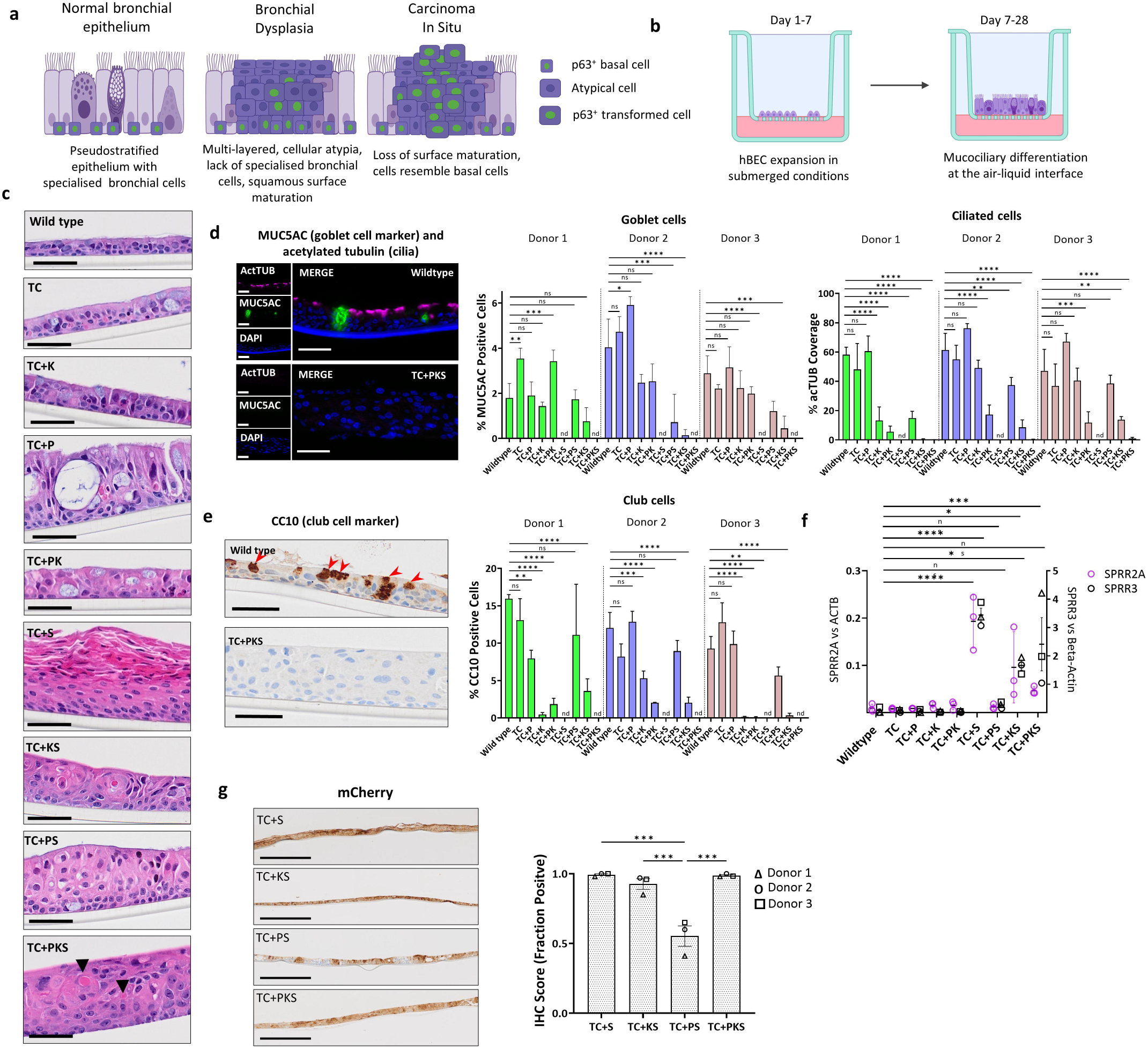
Histological analysis of organotypic ALI cultures generated with mutant hBECs. **a**. Schematic of the developmental stages of LUSC progression. Stages are characterised by reproducible histological changes which include the progression from a mucociliary pseudostratified epithelium to a multi-layered squamous epithelium with loss of differentiation, disordered expression of basal cell markers, loss of surface maturation and cellular atypia. **b**. In vitro air-liquid interface (ALI) organotypic cultures were used to assess histological changes to the airway epithelium driven by LUSC alterations. **c.** Haematoxylin and eosin stained formalin fixed paraffin sections showing the histology of ALI cultures generated using mutant hBECs. Images are representative of three replicate assays carried out using hBECs derived from three donors. Arrowhead indicate areas of keratinisation. Scale bars = 50µm. **d.** Immunofluorescence staining of ALI cultures for MUC5AC (goblet cells, green), acetylated tubulin (cilia, magenta) and DAPI (nuclei, blue) was used to investigate changes in mucociliary differentiation. Left panel shows representative images depicting the staining of one wild type and one TC+PKCS mutant ALI. Right panels show quantification of MUC5AC staining as the % of epithelium composed of goblet cells and acetylated tubulin positivity as the % cilia coverage. Mean +/-SD (n=3). *Adj.P* values were calculated by one-way ANOVA with multiple comparisons and Dunette’s *post hoc* test. nd=not detected Scale bars = 50µm. **e.** Immunohistochemistry staining of ALI cultures for the club cell marker, CC10. Left panel shows images depicting an example of one wild type and one TC+PKCS mutant ALI. Right panel shows quantification of the % CC10 positive cells. Mean +/-SD (n=3). *Adj.P* values were calculated by one-way ANOVA with multiple comparisons and Dunette’s *post hoc* test. nd=not detected. Scale bars = 50µm. **f.** RT-qPCR confirmed the upregulation of genes implicated in squamous differentiation in ALI cultures with SOX2 overexpression. Data is shown as the mean of three independent donors +/-SEM. Adj.P values were calculated by one-way ANOVA with multiple comparisons and Dunette’s post hoc test. ns = not significant. **g**. Immunohistochemistry for the detection of mCherry expression in *SOX2* overexpressing mutant ALI cultures. Left panel shows representative images of mCherry staining from one donor. mutants transduced with the pUltrahot vector carrying *SOX2* cDNA. Right panel shows quantification of mCherry staining as the fraction of epithelium which stained positive. Data is shown as the mean of three independent donors +/-SEM. Donor means were calculated from 3 replicates. *Adj.P* values were calculated by one-way ANOVA with multiple comparisons and Tukey’s *post hoc* test. Scale bars=500µm. ns = not significant.

*SOX2*^OE^ mutants exhibited abrogation of specialized cell differentiation except for TC+PS, which retained higher levels of differentiation (Fig. 3d, e). *SOX2*^OE^ mutants also displayed evidence of squamous differentiation as indicated by apical dyskeratosis, most prominent in TC+S (Fig. 3c and Extended Data Fig. 3a), and expression of the squamous differentiation markers SPRR2A, SPRR3 and IVL (Fig. 3f and Extended Data Fig. 2b). TC+PKS mutants developed keratinisation and a highly disorganised histology with frequent loss of basal-apical polarity, indicative of progression to more advanced developmental stages (Fig. 3c and Extended Data Fig. 2a). We reasoned that lower *SOX2* expression might be responsible for the comparatively abnormal behaviour of TC+PS mutants. Indeed, we found a discontinuous pattern of mCherry expression in TC+PS mutants (Fig. 3g), indicating negative selection of *SOX2*^OE^ cells in TC+PS that is prevented by *KEAP1* inactivation.

LUSC developmental stages are characterised by an expansion of p63-expressing cells from the basal compartment to occupy the epithelial layer. Therefore, we predicted an expansion of p63-positive cells that would be maximal in our modelling of most advanced stages of LUSC. To test this, we first confirmed the expansion of p63^+^ cells in patient specimens with low and high-grade premalignant lesions (Fig. 4a, b). Tissue regions with premalignant lesions were divided into basal and apical halves and the fraction of total p63^+^ cells residing in the apical half was quantified as a measure of p63^+^ cell expansion (Fig. 4a). As expected, we validated an expansion of p63^+^ cells which was maximal in high-grade lesions (Fig. 4b, c). Using the same approach in ALI cultures, we found that p63^+^ cell expansion was maximal in TC+PKS in all three donors (Fig. 4d), strongly indicating that this mutant represents the highest disease grade. Expression of cytokeratins 5 and 6, also a LUSC biomarker, followed a trend similar to p63 (Extended Data Fig. 2b). We also identified inter-donor heterogeneity, most evident in donor 3, where p63^+^ cell expansion was as high in TC+KS as in TC+PKS lacked. This is consistent with the results obtained when investigating anchorage-independent growth in the same mutants from donor 3 (Fig. 2c), which also showed similar levels of colony formation. Of note, expression of the lung adenocarcinoma marker TTF-1 was barely detectable in all mutants (Extended Data Fig. 3a).

**Figure 4.**
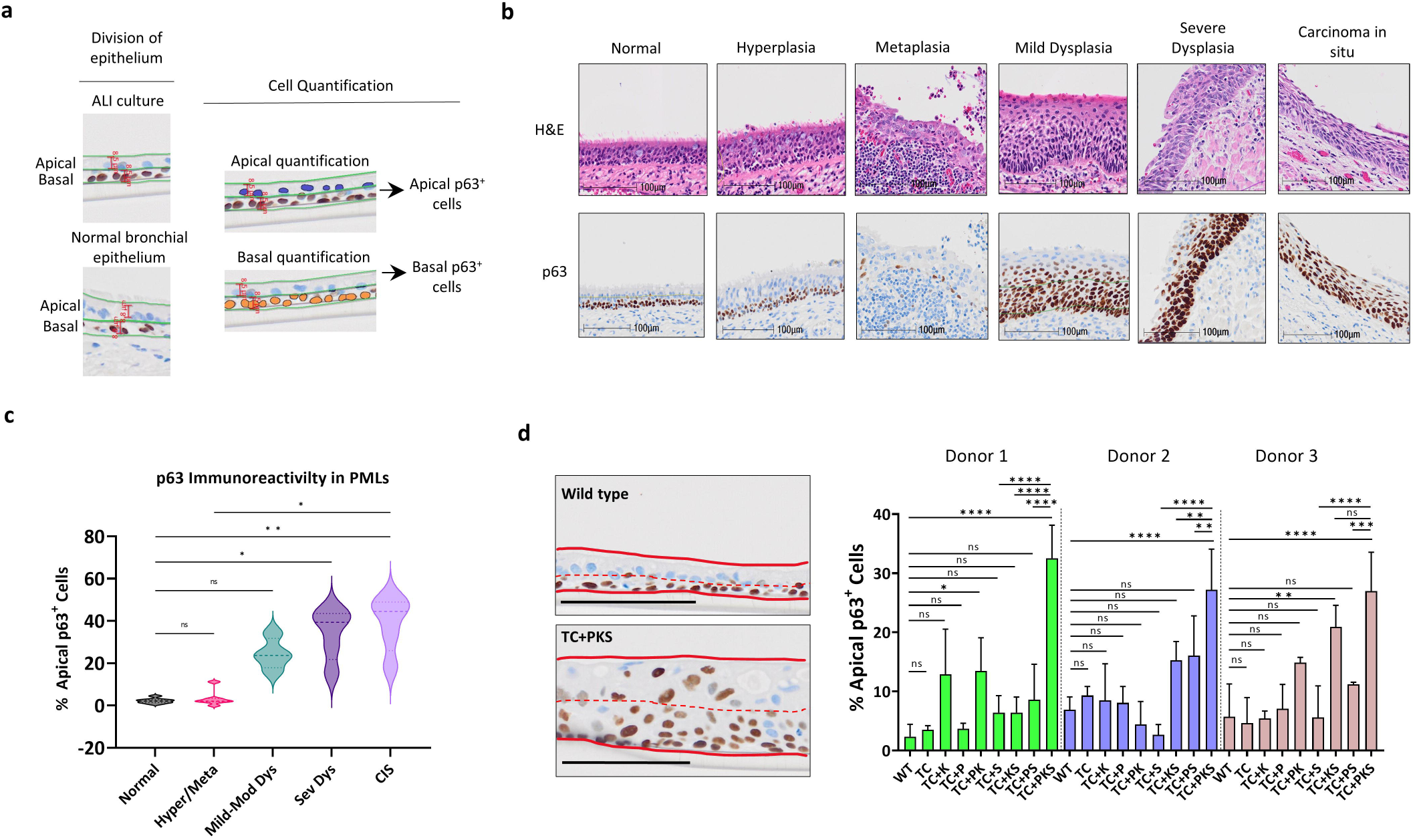
Analysis of p63-expressing cell expansion in LUSC developmental stages and organotypic ALI cultures. **a.** Strategy for the quantification of apical and basal p63^+^ cells in clinical samples (right) and ALI cultures (left). Epitheliums were split laterally into equal halves and separate analyses performed on each half. **b**. Representative haematoxylin and eosin (top) and p63 immunohistochemistry (bottom) staining of LUSC developmental stages in patient clinical samples. **c.** The % of p63 positive cells which reside in the apical half of the airway epithelium is significantly higher in high-grade premalignant lesions than in normal airway epithelium. Analysis used eight clinical samples with different spectrums of premalignant lesions (normal epithelium = 8, hyperplasia/metaplasia = 7, mild-moderate dysplasia = 4, severe dysplasia = 5 and carcinoma in situ = 5). Significance was calculated using Kruskal-Wallis test with multiple comparisons and Dunn’s *post hoc* test. **d.** TC+PKS mutant hBECs generate ALI cultures with a higher proportion of p63^+^ apical cells. Left panel shows representative immunohistochemistry images of wild type and TC+PKS ALI cultures stained for p63. Solid red lines indicate top and bottom of epithelium and dashed red lines mark the lateral division of the epithelium into its apical and basal halves. Scale bars=100µm. The quantification of % p63^+^ apical cells (right panel) in ALI cultures revealed a significant increase in TC+PKS mutation. Mean +/-SD (n=3). *Adj.P* values were calculated by one-way ANOVA with multiple comparisons and Holm-Šídák’s *post hoc* test. ns = not significant.

Taken together, our analyses show that in the absence of lentiviral *SOX2*^OE^, mutant hBECs develop different alterations of the bronchial architecture, but do not acquire clear landmarks of transition to squamous lesions. *SOX2*^OE^ in the TC background induces features of tumour initiation, including multi-layered morphology, acquisition of anchorage-independent growth and squamous differentiation. In TC+PKS mutants, maximal expansion of p63-expressing cells, anchorage-independent growth and invasiveness indicate transition to a malignant phenotype that requires SD, PI3K/Akt and OSR co-operation. Our observations also indicate a detrimental interaction between *SOX2*^OE^ and PI3K/Akt pathway activation that is rescued by OSR pathway activation. This interaction and the requirement of the three pathways for malignant transformation could explain the co-occurrence of 3q-amplification (comprising *SOX2* and *PIK3CA* amplification) and alterations targeting the OSR pathway in the classical LUSC subtype. In line with our results, interrogation of DepMap Consortium data (https://depmap.org/portal/) (16, 17) revealed that LUSC was the most dependent on *NFE2L2* of all tumour types (Extended Data Fig. 3b, c). *NFE2L2* encodes for Nrf2, a master regulator of the OSR (or Nrf2) pathway that is regulated by *KEAP1* (Extended Data Fig. 3d). *SOX2* is also a co-dependency in *NFE2L2*-dependent cell lines, implying that *NFE2L2*-dependent cell lines are also dependent on *SOX2* (Extended Data Fig. 3e, f).

### Comprehensive gene-expression analysis of dysregulated pathways

Our combinatorial LUSC modelling approach allows us to deconvolve the effect of pathway dysregulation on gene-expression. To do this, we carried out bulk RNA-seq of ALI cultures. Before focusing on the effect of specific pathways, we carried out PCA and WGCNA analyses to confirm that mutants from different donors behave similarly at the gene expression level. PCA revealed four clusters segregated primarily by genotype (*KEAP1* loss and *SOX2*^OE^) instead of donor (Extended Data Fig. 4a). As expected, TC+PS consistently clustered with wild type, TC and TC+P mutants. The role of *SOX2* and *KEAP1* in shaping LUSC transcriptomes is congruent with published studies (4, 8, 10). Next, WGCNA was performed to assess inter-donor heterogeneity (Extended Data Fig. 4b, c) (18). Seven consensus co-expression modules were identified when gene expression from the three donors was analysed globally (Extended Data Fig. 4b, Supplementary Tables 3, 4). These seven modules were largely conserved amongst the three donors and only a few additional small modules with low stability were identified in individual modules (Extended Data Fig. 4c, 5a-d). After identifying limited heterogeneity between donors, we set out to analyse the transcriptomic data globally and identify differentially expressed genes regulated by each pathway relative to the TC background (Supplementary Table 5) followed by gene-set enrichment analysis (GSEA) (Fig. 5a). TC+PKS relative to TC were also interrogated to ensure gene-set enrichments attributed to one pathway is not supressed by the concomitant dysregulation of additional pathways (Figure 5a, Supplementary Table 5, 6). Due to the size of the transcriptomic data, we focused on GO Biological Process and Hallmark collections. Positively enriched BP GO terms associated with *SOX2*^OE^ were dominated by epithelial development and keratinocyte biology (Fig. 5b and Supplementary Tables 5, 7), reflecting transition to a squamous keratinised epithelium, characteristic of squamous metaplasia. Genes upregulated in TC+S and TC+PKS mutants were also enriched in RAS-RTK pathway associated GO terms (Fig. 5b, and Supplementary Table 5, 7) and in the KRAS SIGNALING UP hallmark (Fig. 5c, Extended Data Fig. 6a and Supplementary Table 6, 7). RAS-RTK signalling in LUSC has been functionally linked to *SOX2* in cooperation with *TP63* to induce *EGFR* expression (19) and with the expression of EGFR ligands different from EGF (8). We confirmed that *SOX2*^OE^ upregulated *TP63* and EGFR ligands, but not EGFR. (Fig. 5d and Supplementary Tables 5). We identified a heterogeneous group of positively enriched GO terms related to protein processing, including multiple serine protease inhibitors (Fig. 5b, e, Extended Data Fig. 6b and Supplementary Table 5, 7). Among these was *PI3,* which encodes a neutrophil-elastase inhibitor and known component of the cornified envelope (20), highly upregulated in *SOX2*^OE^ mutants (Extended Data Fig. 6c, Supplementary Table 5). Therefore, we hypothesised that *PI3* might counteract the known tumour suppressive activity of neutrophil-elastases to promote LUSC progression (21). A significant positive correlation between *SOX2* and *PI3* mRNA (Spearman=0.24 q-value=0.0335) and protein (Spearman=0.268, q-value=0.0214) expression was confirmed using CPTAC data (8), but not the TCGA cohort (9). When we interrogated a transcriptomic database of premalignant lesions previously published (22, 23), maximal *PI3* expression was observed in high-grade and invasive carcinomas (Fig. 5f), implying a function in the transition to invasive carcinomas.

**Figure 5.**
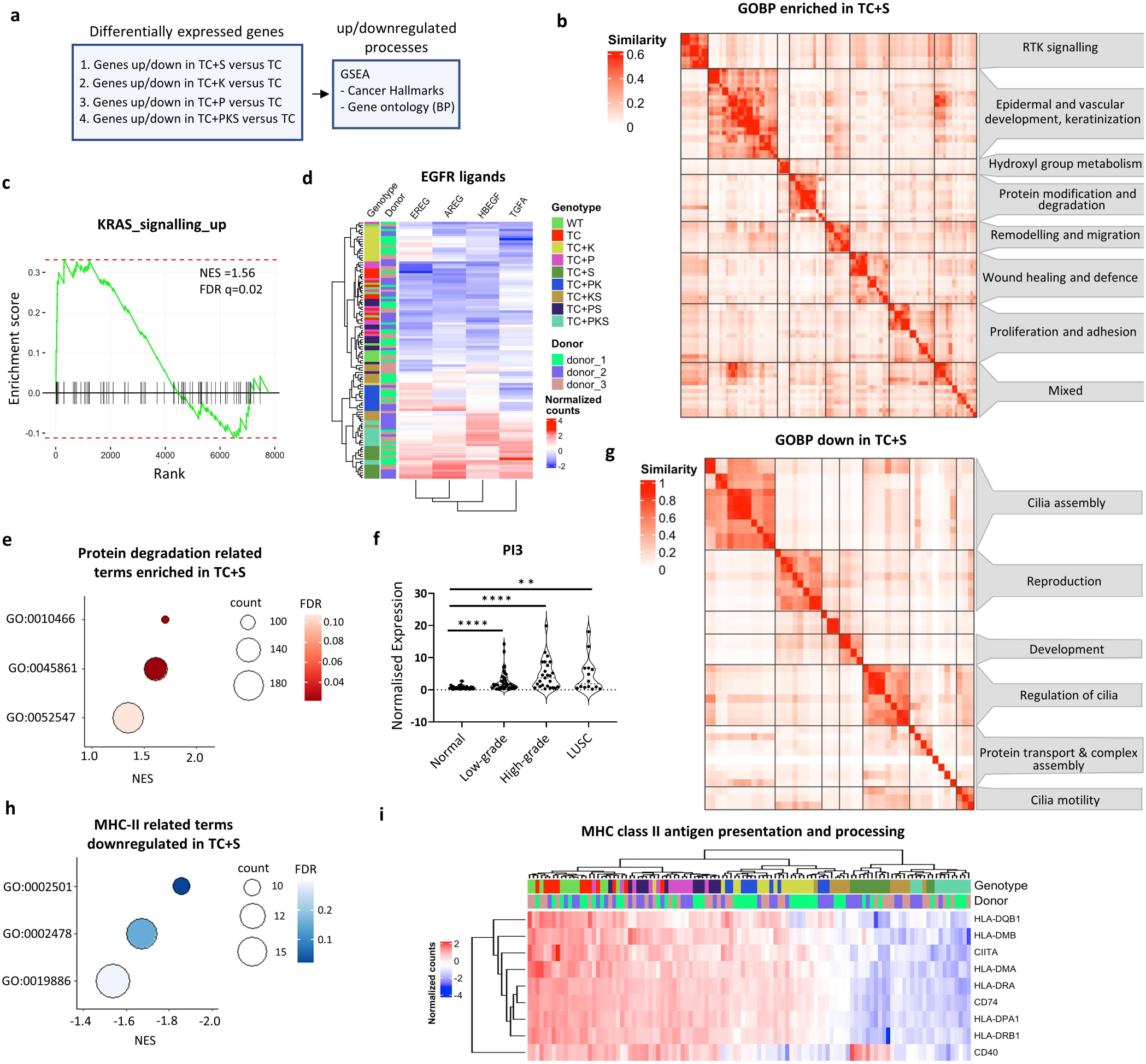
Global transcriptomic effects of *SOX2*^OE^ in mutant hBECs. **a.** Strategy for the identification of processes associated with each single pathway (TC+S/K/P) and all pathways in combination (TC+PKS). Gene expression changes associated with the dysregulation of each single pathway and all pathways in combination were identified by carrying out differential gene expression (TC+P/S/K/PKS versus TC). Gene set enrichment analysis (GSEA) was performed with differentially expressed genes. **b.** Clustergram of significant positively enriched gene ontologies (Biological Process) using gene set enrichment analysis of genes differentially expressed in TC+S versus TC comparisons (q-value≤ 0.05). Terms were aggregated using a measure of semantic similarity (similarity) using the simplifyEnrichment package. **c**. Gene set enrichment analysis showing the significant positive enrichment (q-value<0.05) of the KRAS signalling up hallmark (MSigDB, H: hallmark gene sets) in genes differentially expressed in TC+S versus TC (q-value ≤ 0.05). **d.** Heatmap and hierarchical clustering showing the normalised expression of EGFR ligands across the entire sample set. **e.** Dot plot showing positively enriched gene ontology terms related to protein degradation within the ‘protein modification and degradation’ cluster from (b). GO:0010466=negative regulation of peptidase activity, GO:0045861=negative regulation of proteolysis, GO:0052547=regulation of peptidase activity. **f**. Violin plot showing the expression of *PI3* in normal (n=42), low (n=41) and high-grade (n=25) premalignant lesions and lung squamous cell carcinoma (n=14). Significance was calculated by comparing the means of each group to ‘normal’ using the Mann-Whitney test. GEO accession: GSE33479. **g.** Clustergram of significant (q-value<0.05) negatively enriched gene ontologies (GO Biological Process) using gene set enrichment analysis of genes differentially expressed in TC+S versus TC comparisons (q-value ≤ 0.05). **h.** Dot plot showing negatively enriched gene ontology terms related to MHC-II peptide presentation within the ‘protein transport and complex assembly’ cluster from (g). GO:0019886=antigen processing and presentation of exogenous peptide antigen via MHC class II, GO:0002478=antigen processing and presentation of endogenous peptide antigen, and GO:0002501=peptide antigen assembly with MHC protein complex. **i**. Heatmap and hierarchical clustering showing the normalised expression of gene involved in MHC-II antigen processing and presentation across the entire sample set.

As expected, processes downregulated following *SOX2*^OE^ largely related to cilia biology (Fig. 5g and Supplementary Tables 5-7), reflecting bronchial-squamous reprogramming (Fig. 3c-f). A cluster of negatively enriched ontologies related to protein transport contained gene-sets consisting of MHC-II subunits (Fig. 5g-i, Extended Data Fig. 6d and Supplementary Table 5-7). Two recently published articles that analyse premalignant lesions (24) or use LUSC human and mouse models (25) reported the downregulation of epithelial MHC-II expression mediated by the MHC-II transcriptional activator, *CIITA*. However, our data showed that *CIITA* was downregulated by *SOX2*^OE^ (Fig. 5i and Supplementary Table 5). Furthermore, we confirmed a negative correlation between *CIITA* expression and MHC-II genes in LUSC with *SOX2* expression using both the TCGA and CPTAC cohorts (Extended Data Fig. 6e, f). Our analysis of SOX2-associated expression data strongly indicates that *SOX2* amplification in LUSC is multi-faceted and constitutes a hub of pro-tumourigenic cues, not limited to cell autonomous processes (squamous differentiation, RTK-RAS pathway activation), but also to remodel the immune microenvironment.

Consistent with the function of the OSR pathway, *KEAP1* inactivation resulted in positive enrichment of redox metabolism and xenobiotic detoxification gene-sets (Fig. 6a, and Supplementary Table 8). A similar set of significantly enriched terms has been identified in the classical LUSC subtype, highlighting the importance of this pathway in shaping LUSC transcriptomes (4, 8, 10). Further to this, we observed an enrichment of amino acid transport gene-sets known to be bona-fide Nrf-2 targets (Fig. 6a-c, Extended Data Fig. 7a and Supplementary Table 5, 6 and 8). We confirmed upregulation of a subset of these transporters in LUSC tumours with OSR-targeting alterations; *SLC1A4* and *SLC1A5* (glutamine influx), *SLC7A11* (cystin influx), SCL75A (essential amino acids), LRRC8D (anion channel) and *SLC3A2 (*ancillary to *SLC7A5* and *SLC7A11)* (Fig. 6d). To the best of our knowledge, *LRRC8D* has not previously been reported as an Nrf-2 target but has been linked to chemotherapy resistance (24). Finally, we observed an enrichment in the *glucose-6-phosphate metabolic process* GO term (lipid and glucose metabolism cluster) (Fig. 6a, e, and Supplementary Table 5, 6 and 8) that included most pentose-phosphate pathway (PPP) enzymes. These enzymes are known Nrf-2 targets and are upregulated in LUSC with OSR alterations (Fig. 6f). PPP upregulation via OSR activation contributes to redox homeostasis by maintaining high GSH/GSSG ratios and NADPH production, although other redox metabolism unrelated functions, such as nucleotide biosynthesis, could play an important role.

**Figure 6:**
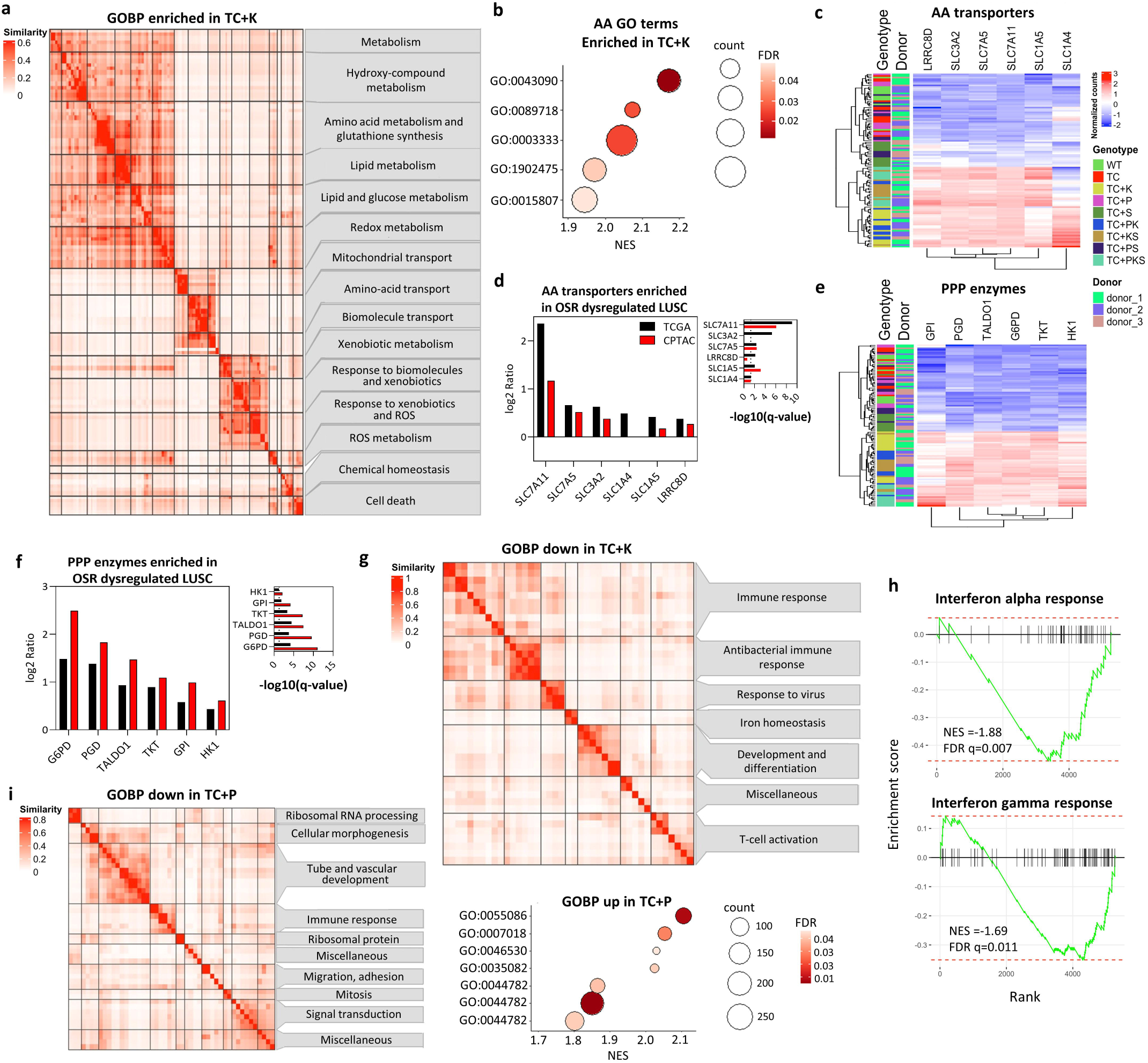
Global transcriptomic effects of *KEAP1* and *PTEN* inactivation in mutant hBECs. **a.** Clustergram of significant positively enriched gene ontologies (GO Biological Process) using gene set enrichment analysis of genes differentially expressed in TC+K versus TC comparisons (q-value< 0.05). **b.** Dot plot showing positively enriched gene ontology terms related to amino acid transport within the ‘ammino acid transport’ cluster from (a). GO:0015807=amino acid transport, GO:0003333= amino acid transmembrane transport, GO:1902475=alpha amino acid transmembrane transport, GO:0089718=amino acid import across plasma membrane, and GO:0043090=amino acid import. **c.** Heatmap and hierarchical clustering showing the normalised expression of amino acid transporters across the entire sample set. **d.** Bar chart showing Log2Ratios comparing the expression of amino acid transporters between LUSC tumours with and without NRF2 pathway alterations (KEAP1, NFE2L2, and CUL3) in 511 LUSC samples from the TCGA (mRNA) and 80 LUSC samples from CPTAC (protein). Left panel shows Log2Ratios. Right panel shows -log10(q-value). Dotted line indicates significance threshold (q-value<0.05). p-values were derived from student’s t-tests and q-values from Benjamini-Hochberg procedure. **e.** Heatmap and hierarchical clustering showing the normalised expression of pentose phosphate pathway (PPP) enzymes in the entire sample set. **f.** Bar chart showing Log2Ratios comparing the expression of pentose phosphate pathway (PPP) enzymes between LUSC tumours with and without NRF2 pathway alterations (*KEAP1*, *NFE2L2*, and *CUL3*) in 511 LUSC samples from the TCGA (mRNA) and 80 LUSC samples from CPTAC (protein). Left panel shows Log2Ratios. Right panel shows-log10(q-value). Dotted line indicates significance threshold (q-value<0.05). p-values were derived from student’s t-tests and q-values from Benjamini-Hochberg. **g.** Clustergram of significant (q-value<0.05) negatively enriched gene ontologies (GO Biological Process) using gene set enrichment analysis of genes differentially expressed in TC+K versus TC comparisons (q-value ≤ 0.05). **h.** Gene set enrichment analysis showing the significant negative enrichment (q-value<0.05) of the response to interferon alpha and (top) and response to interferon gamma (bottom) hallmarks (MSigDB, H: hallmark gene sets) in genes differentially expressed in TC+K versus TC (q-value < 0.05). **i.** Clustergrams of negatively and positively enriched gene ontologies (GO Biological Process) (FDR<0.05) in gene set enrichment analysis comparing TC+P versus TC (q-value ≤ 0.05, log2FC).

*KEAP1* inactivation drove the downregulation of immunity-related gene-sets (Fig. 6g and Supplementary Table 8) possibly reflecting the negative enrichment for interferon response hallmarks (Fig. 6h and Supplementary Table 8) and the enrichment of both hallmarks that we observed in module 6 genes (downregulated *KEAP1* mutants) of our WGCNA analysis (Extended Data Fig. 4b and 7b). This immuno-suppressive role through interferon response inhibition by OSR has been reported in lung adenocarcinoma but not in LUSC (25). Whilst TC+PKS mutants showed similar results, they did not reach significance for the interferon gamma response hallmark (Extended Data Fig. 7c). Likewise, tumours with OSR-targeting alterations also showed downregulation of genes in interferon and inflammation related hallmarks (Extended Data Fig. 7d, Supplementary Table 4), and suppression of interferon responses was most prominent in the classical subtype (8). Our model therefore recapitulates a wide range of reported metabolic and immune-related transcriptional changes mediated by OSR activation.

PI3K/Akt activation by *PTEN* truncation resulted in a more limited transcriptional response than *SOX2*^OE^ and *KEAP1* truncation (Extended Data Fig. 7e and Supplementary Table 5). This is consistent with our PCA and WGCNA analysis (Extended Data Fig. 4a, b) showing that the latter alterations are the most determinant factors in shaping transcriptomes, and on the other hand, with reports showing that LUSC subtypes are not determined by PI3K/Akt-targeting alterations (8, 10). GSEA analysis showed the expected positive enrichment in the MTORC1 and Glycolysis hallmarks (Extended Data Fig. 7e, f, Supplementary Table 9) and predominantly negative enrichment in GO terms related to multiple processes, such as ribosome biology, migration and development (Fig. 6i, j, Supplementary Table 9).

### An evolutionary sequence of pathway activation in LUSC development

Although the main aim of our combinatorial strategy of genetic manipulation was to unravel how the most relevant LUSC pathways cooperate to drive LUSC progression, our results can be mined to hypothesize the most likely trajectory of pathway activation. Our morphological analysis revealed that *SOX2*^OE^ abrogates bronchial epithelial architecture and induces SD (Fig. 3c-e, and Extended Data Fig. 7a, b), two processes that occur early in LUSC development, whereas additional *KEAP1* and *PTEN* inactivation drive further p63-positive cell expansion and invasiveness (Fig. 2d and 4d). Therefore, *SOX2* amplification, or more generally, activation of the SD pathway, might occur earlier than OSR and PI3K/Akt pathways (Fig. 7a). Additionally, our observation that *KEAP1*-loss rescues the negative interaction between *SOX2*^OE^ and *PTEN* truncation suggests that PI3K/Akt pathway activation might occur at an evolutionary stage that avoids the detrimental interaction with SOX2 (Fig. 7a). Therefore, according to our results, the sequence SD-OSR-PI3K/Akt is the most likely evolution of pathway activation according to our results (Fig. 7a). We next ascertained whether this hypothetical evolutionary trajectory is consistent with observations in human specimens. Published LUSC evolutionary histories (26) lack the temporal resolution required for this analysis. Moreover, the lack genomic characterisations of developmental LUSC stages prevents us from inferring an evolutionary sequence of somatic events. However, the preinvasive lesion cohort generated by Mascaux and colleagues (22) can be interrogated to map the onset of pathway dysregulation using transcriptional signatures. Thus, we carried out ssGSEA analysis on the latter data using gene-sets associated with the three pathways (Fig. 7b-d). As expected, an increase in squamous differentiation-related enrichment scores was significant in squamous metaplasia and increased OSR- and PI3K/Akt-related scores became significant in squamous metaplasia and moderate dysplasia respectively (Figure 7c, d). These observations support that PI3K/Akt pathway activation occurs later than SD and OSR pathways, which appear to be upregulated simultaneously. Though stromal contamination, LUSC molecular subtypes and the assumption that LUSC development is homogeneous across patients might be confounding factors, these results indicate that the onset of PI3K/Akt signalling occurs later in LUSC evolution, perhaps due to a reduction in cell fitness caused by the interaction between PI3K/Akt and *SOX2* gain-of-function.

**Figure 7:**
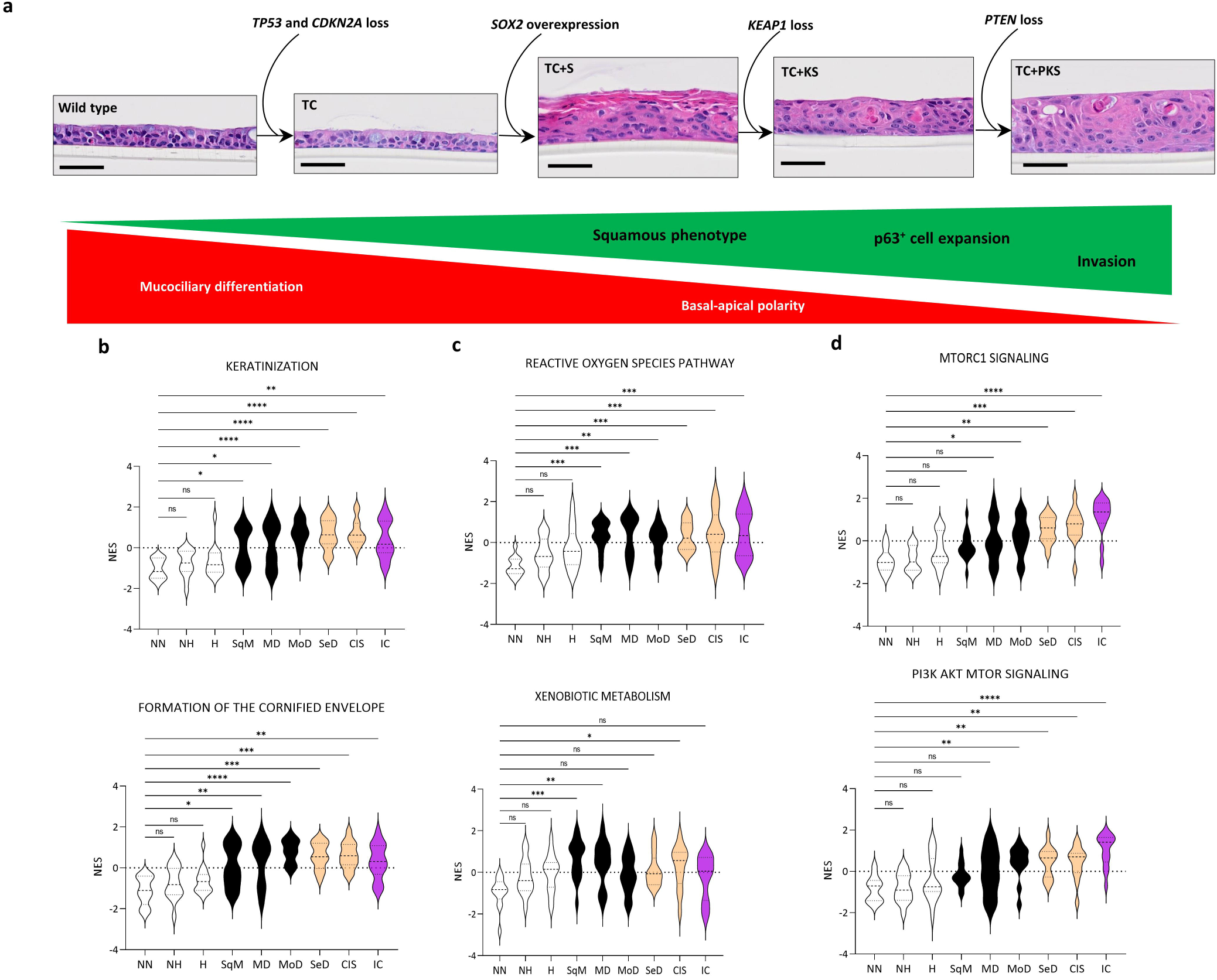
A hypothetical evolutionary sequence of pathway activation in LUSC development. **a**. Hypothesised evolutionary sequence of pathway activation in LUSC development based on the morphological and molecular characteristics of our model including loss of mucociliary differentiation and epithelial polarity, and gain of a squamous phenotype, expansion of p63^+^ cells, and acquisition of an invasive phenotype. Scale bars = 50µm. **b-d**. Normalised enrichment scores (ssGSEA) calculated from the dataset published by Mascaux et al., (2019) (GEO accession: GSE33479) for gene signatures associated with activation of the squamous differentiation pathway (**b**) including keratinisation (GO:0031424, top) and formation of the cornified envelope (Reactome R-HSA-68009371, bottom), the OSR pathway (**c**) including the reactive oxygen species pathway (Hallmark M5938, top) and xenobiotic metabolism (Hallmarks M5934, bottom), and the PI3K/AKT pathway (**d**) including MTOCR1 signalling (Hallmarks, M5924, top) and PI3K AKT MTOR signalling (Hallmarks M5923, bottom). Scores are shown for each stage of LUSC development including normal (NN), normal hypofluorescent (NH), hyperplastic (H), squamous metaplasia (SqM), mild dysplasia (MD), moderate dysplasia (MoD), severe dysplasia (SeD), carcinoma in situ (CIS), lung squamous cell carcinoma (LUSC). White = normal, black = low grade, green = high grade, and red = invasive carcinoma. Significance (q-value<0.05) was calculated using Kruskal-Wallis testing with multiple comparison and Dunn’s *post hoc* test.

## DISCUSSION

Advances in translational cancer research rely on the availability of tractable models that recapitulate key cancer traits. Mouse models are valuable for understanding basic LUSC biology. However, availability of resources and inter-species differences hinder the modelling of increasingly complex genotypes, evolutionary trajectories and inter-patient heterogeneity. We demonstrated here that genetically engineered hBECs constitute a tractable, biologically relevant system to study LUSC as demonstrated by the expression of p63 and cytokeratins 5/6, absence of TTF-1, and presence of keratinisation. We implemented our strategy with the same cell type from three donors without cancer or airway disease to ensure reproducibility. Similar models developed by genetic manipulation of primary human cells published for other cancer types made use of only one donor in a melanoma model (27), or cells isolated from multiple patients with colorectal cancer (28) and cells from spatially separated intestinal regions (29) in colorectal cancer models. Overall, our cell biology results indicated that concomitant activation of the SD, PI3K/Akt and OSR pathways was required for malignant transformation. Although some inter-donor heterogeneity was detected, a holistic vision of our results revealed homogeneous phenotypes in the three donors who were of similar age, non-smokers and lacked airway disease.

We conclude that our approach has generated a human model of the classical LUSC subtype (4, 8) as [1] the three pivotal pathways investigated were required for invasive transformation which is consistent with the accumulation of OSR pathway mutations and 3q-amplification (*SOX2* and *PIK3CA*) in classical LUSC subtype (8), [2] OSR activation was required for the expansion of *SOX2*^OE^ cells with concomitant PI3K/Akt activation, and [3] OSR and *SOX2* downregulated pathways involved in tumour immunity, which could contribute to the ‘immune-cold’ environment observed in the classical LUSC subtype. Mouse models have not clearly demonstrated the requirement for SD, PI3K/Akt and OSR activation in defining classical LUSC subtype. Ferone et al., (2017) used a strategy that involved *Sox2*^OE^and *Pten*^ko^ without targeting the OSR pathway that successfully resulted in tumourigenesis. Notwithstanding inter-species differences and the lack of evidence of invasion in the latter model, positive selection for cells with OSR activation during tumour development in this model might explain this difference.

Our observations suggest new translational perspectives. For example, our results indicate that tumours with 3q-amplification and OSR-targeting alterations are likely to be more sensitive to experimental therapies targeting the Nrf-2 pathway (e.g. glutaminase inhibitors) and that rescuing the negative interaction between SOX2 and PI3K/Akt activation might constitute an elusive basic function of OSR activation in LUSC. A likely mechanism could be OSR mediated detoxification of endogenous toxic ROS originating from cumulative *SOX2* and PI3K/Akt driven metabolic reprogramming (30, 31) or increased metabolic demands caused by concomitant *SOX2*^OE^ and PI3K/Akt, such as a higher demand for amino acids and glutaminolysis reliance, that are compensated by upregulation of essential amino acid (*SLC7A5*) and glutamine (*SLC1A5*) transporters. In fact, our results might explain why lung cancer prevention strategies based on the use of antioxidants actually increase the risk of lung cancer, as these agents mimic Nrf-2 pathway activation and might favour the expansion of clones with 3q-amplification (30, 31).

Therapies that target the OSR pathway by limiting glutamate biosynthesis for glutathione production have been tested in trials (NCT04265534 and NCT04471415) without positive result. Addressing this lack therapeutic effect and testing new strategies to target the OSR pathway can be approached with our human model as it has recapitulated the wide range of gene expression changes previously reported for the OSR pathway, and additional changes, reflecting a complex interplay between interdependent and/or functionally redundant amino acid transporters.

We also show regulation of innate and adaptive immunity by the SD and OSR pathways. SOX2 downregulates the neutrophil-elastase inhibitor *PI3* and MHC-II while OSR activation lowered type-I and II interferon responses. For the first time, we have provided evidence that places SOX2 at the apex of cell intrinsic and TME-mediated pro-oncogenic signals that include links with innate and acquired immunity in LUSC such as MHC-II downregulation and neutrophil-elastase inhibitors. Notwithstanding that potential strategies to target the individual processes mediated by SOX2 must be tested (21, 25), the multi-faceted SOX2 oncogenic functions emphasise the need to target SOX2 itself. New therapeutic tools such as PROTACs or approaches analogous to Omomyc could facilitate this endeavour (32). Similarly, OSR inhibition is likely to enhance type-I and II interferon responses and remodel the immune cell landscape of LUSC. This is in line with the limited therapeutic response to immunotherapies observed in cancer patients and mouse models with OSR alterations (25, 33, 34). Our human LUSC model also provide a new experimental tool to investigate the latter strategies.

Finally, the comparison of our observations with transcriptomic signatures in patient samples are in accordance with an evolutionary trajectory in which activation of SD and OSR programs precede PI3K/Akt activation, but with several caveats. This trajectory is at odds with *SOX2* and *PIK3CA* co-amplification in 3q26, which suggests simultaneous SD and PI3K/Akt activation. However, gene-dose to phenotype correlations likely differ amongst genes, as might the timing of phenotype emergence following a copy number event. In fact, 3q26 amplification is detectable in progressive squamous metaplasias (35) and increases during LUSC development (36). If *PIK3CA* gene dose is less efficient in inducing the pathway, this would result in an evolutionary ‘delay’ of PI3K/Akt activation.

We set out to provide an informative, tractable *in vitro* LUSC model as an alternative, or an animal use reducing prelude, to more expensive, technically challenging and animal consuming *in vivo* modelling of LUSC. By definition, our model focuses primarily on cell-intrinsic effects lacking the complexity of the native TME, which constitutes a limitation. However, our model incorporates intra-epithelial conditions and can be progressively upgraded to incorporate TME components. Our model can now be applied to address further basic and translational research questions, the most obvious being a reverse genetic approach to model LUSC heterogeneity using comparative genomic studies. For instance, our system can be used to test whether other drivers of the three LUSC prevalent pathways phenocopy the alterations used in this study and trigger the same vulnerabilities or, alternatively, to assess the function of unrelated drivers. Our model also provides an opportunity to explore the biology of risk factors including smoking and COPD by manipulating hBECs derived from appropriate donors. Although the study of metastasis is an additional limitation of our model, the use of bioengineering solutions, such as the organ-on-a-chip technology, might overcome this limitation. Lastly, identification of new biomarkers for LUSC prevention and early detection methods, drug resistance, and immune-oncology studies are examples of the multiple translational opportunities to exploit in our new LUSC model.

## AUTHOR CONTRIBUTIONS

JO: Conceptualization, data curation, formal analysis, investigation, methodology, validation, visualization and writing-original draft, review & editing

RS: Conceptualization, data curation, formal analysis, methodology, software, validation

SS: Conceptualization, data curation, formal analysis, methodology, software, supervision and validation

AO: Data curation, investigation, methodology, project administration, resources, validation and writing-review & editing

AC: Data curation, investigation and resources

CD: Conceptualization, funding acquisition, methodology, resources, supervision and writing-review & editing

CLG: Conceptualization, data curation, formal analysis, funding acquisition, methodology, project administration, resources, supervision, validation, visualization and writing-original draft, review & editing

## ACKNOWLEDGEMENTS

This project has been funded by the National Centre for the Replacement, Refinement & Reduction of Animals in Research (NC/W001284/1), the Rosetrees Trust (M767), the Cancer Research UK Lung Cancer Centre of Excellence (A25146) and the Cancer Research UK Manchester Institute (C5759/A20971).

We would like to thank the Molecular Biology, Histopathology, Flow Cytometry and Scientific Computing core facilities at the Cancer Research UK Manchester Institute, the Cancer Biomarker Centre, and the Manchester Cancer Research Centre Biobank.

Patient samples were obtained from the Manchester Cancer Research Centre (MCRC) Biobank (Approval #23_CALO_01). The MCRC Biobank holds a generic ethics approval (Ref: 18/NW/0092) which can confer this approval to users of banked samples via the MCRC Biobank Access Policy.

## EXTENDED DATA FIGURE LEGENDS

**Extended Data Figure1.**
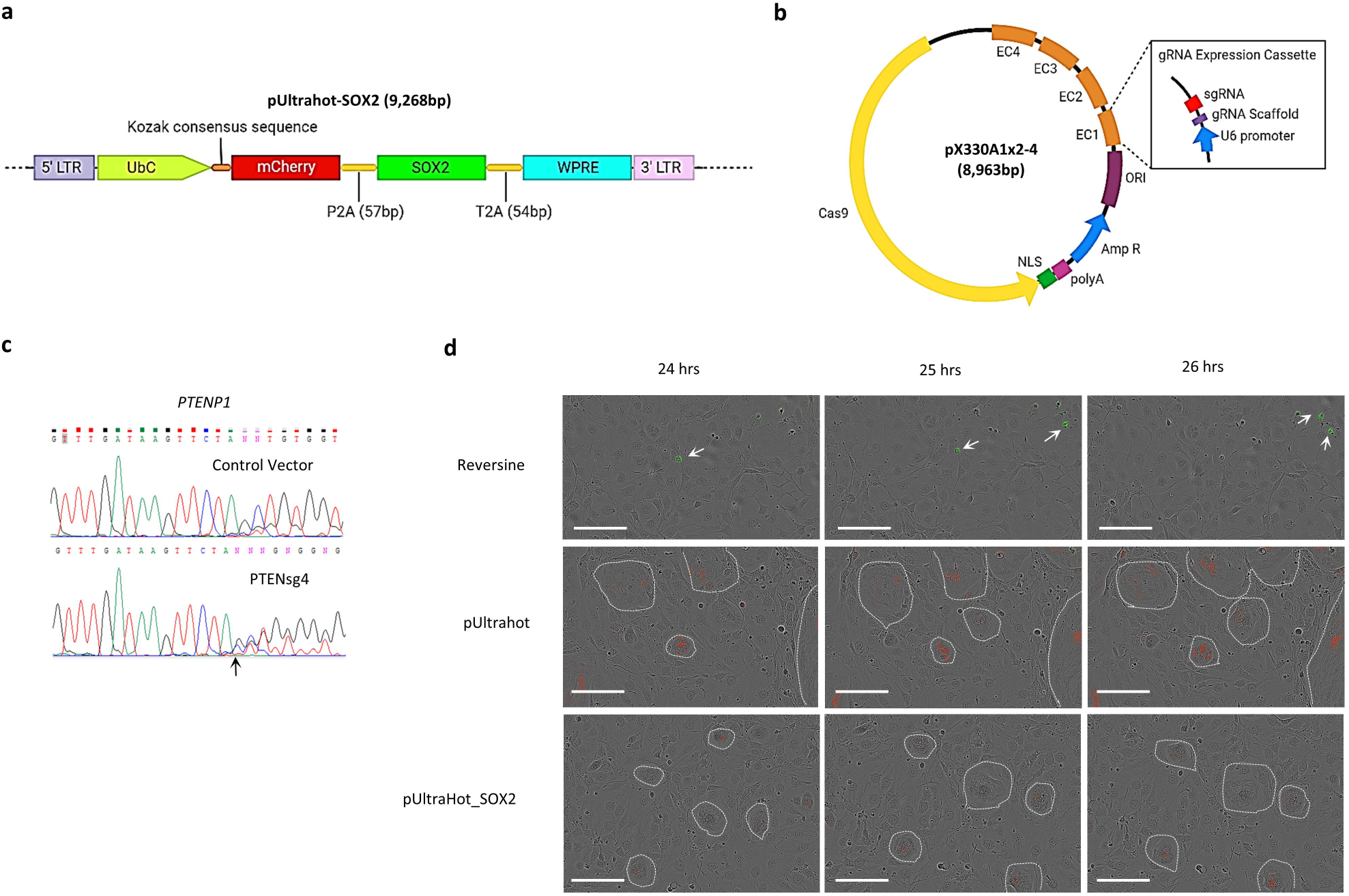
**a**. Diagram of pUltrahot construct insert region with mCherry reporter and SOX2 cDNA intersected by a P2A self-cleaving peptide sequence. **b**. Diagram of the pX330 multiplex CRISPR/Cas9 vector used to generate tumour suppressor knockouts. The vectors contained a maximum of four gRNA sequences inserted into individual gRNA expression cassettes (EC). **c**. Sanger sequencing traces of the *PTENP1* locus from hBECs electroporated with the empty pX330 CRISPR/cas9 vector (control vector) and those electroporated with a px330 CRISPR/cas9 vector containing the PTEN gRNA #4. **d**. Representative images of cleaved caspase 3 apoptosis assays depicting live cell imaging of hBEC cultures transduced with the empty pUltrahot vector, or pUltrahot with SOX2 cDNA (pUltrahot_SOX2). Imaging was carried out every 60 minutes from timepoints 0 to 96-hours. Images show 24-, 25- and 26-hour timepoints. hBECs treated with reversine acted as an apoptosis positive control. hBEC colonies are identified by dashed white lines and mCherry expressing hBECs are shown with red fluorescence. Cells undergoing apoptosis appear in green and are highlighted by white arrows. Scale bars = 200µm.

**Extended Data Figure 2.**
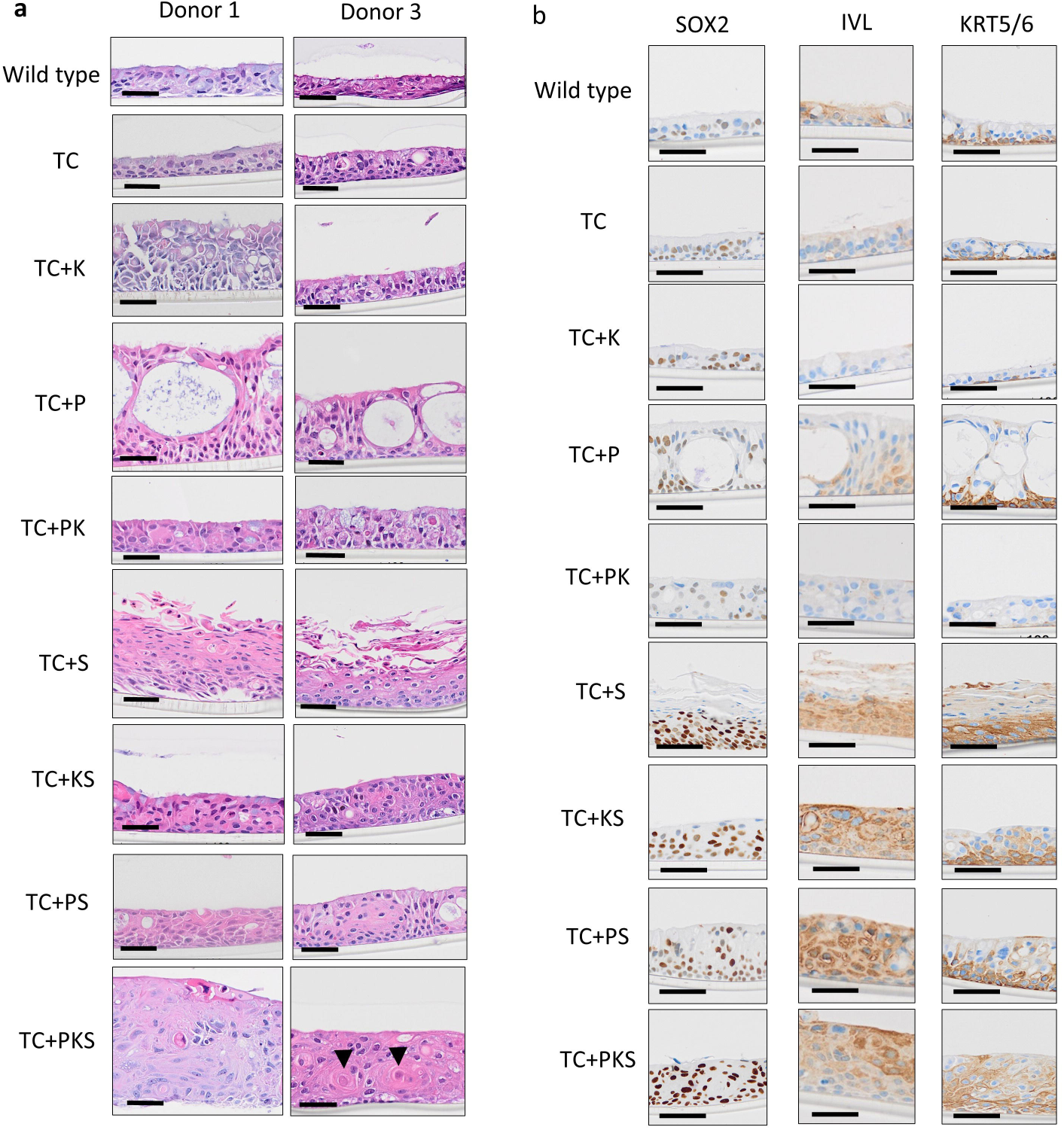
**a.** Representative haematoxylin and eosin stained formalin fixed paraffin sections showing the histology of ALI cultures generated using mutant hBECs from donor 1 and 3. Arrowheads show areas of keratinisation. Scale bars = 50µm. **b**. Representative images of **i**mmunohistochemistry staining of ALI culture formalin fixed paraffin embedded sections to detect the expression of SOX2 and proteins associated with terminal squamous differentiation in donor 2. IVL = involucrin, KRT5/6 = cytokeratin 5/6. Scale bars = 50µm.

**Extended Data Figure 3.**
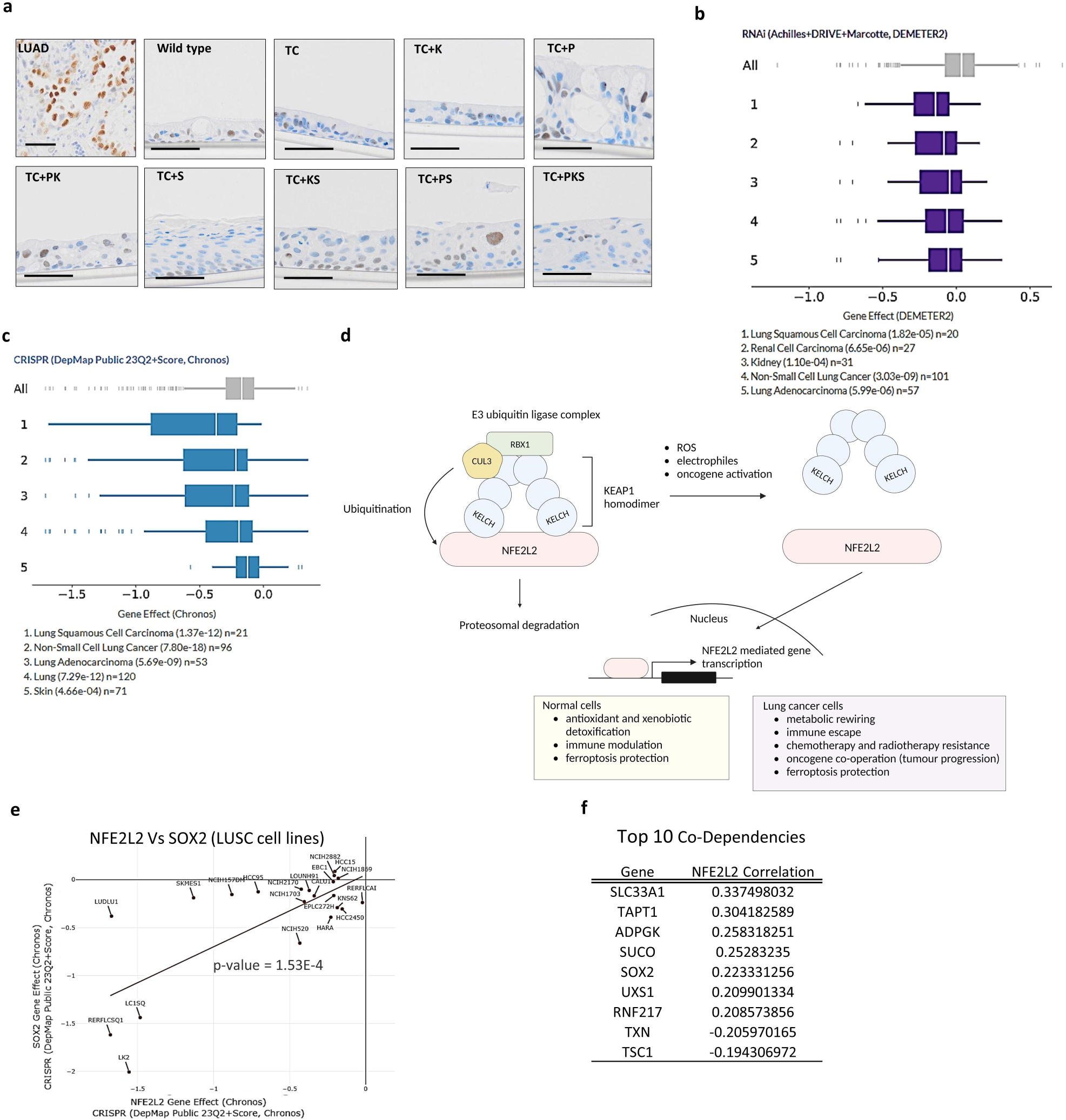
**a.** Immunohistochemistry staining of ALI cultures for the pulmonary developmental transcription factor TTF-1. TTF-1 is positive in LUAD control samples but expressed at low levels in ALI cultures. **b-c.** DepMap data showing cell line lineages ranked *NFE2L2* gene effect in both the RNAi (**b**) and CRISPR (**c**) datasets. Top 5 cancer types are shown. **d.** DepMap co-dependency data showing the top 10 significant genes correlated with *NFE2L2* expression. Genes are ranked by correlation coefficient. **e**. DepMap data showing the correlation between *NFE2L2* and *SOX2* gene effects in LUSC cell lines. *NFE2L2* and *SOX2* gene effects are significantly correlated (p-value = 1.53E-4). **f.** A schematic of the KEAP1-NRF2 pathway with examples of functional outputs in normal and cancer context.

**Extended Data Figure 4.**
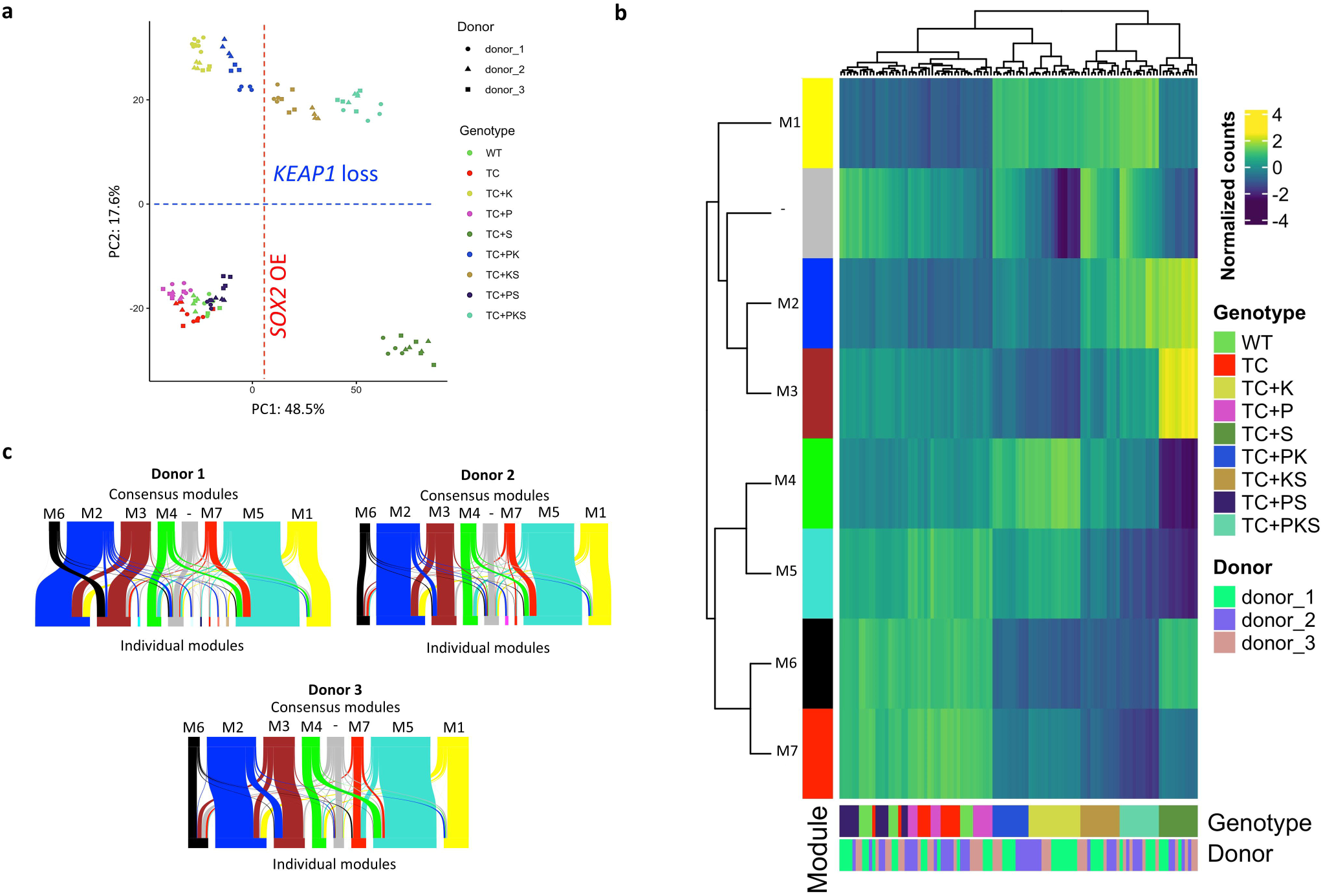
**a**. Principal component analysis (PCA) using the top 500 variable genes from RNAseq of 108 ALI cultures. Four ALI cultures from each genotype were included, each from one of the of three hBEC donors. PC1 separated samples based on the presence or absence of *SOX2* (red dotted line) and PC1 separated samples based on the presence or absence of functional *KEAP1* (blue dotted line). **b** Seven consensus modules of co-expressed genes were identified using weighted gene co-expression network analysis (WGCNA). Modules are labelled M1-M7. Genes marked by the grey bar did not fit any specific co-expression module. Heatmap indicates the average normalised counts of module gene sets within each sample. Hierarchical clustering was performed based on both genotype and module expression. **c.** Sankey diagrams showing the relationship between the seven consensus modules (identified using all data) and the individual donor modules (identified separately for each donor) for donors 1-3.

**Extended Data Figure 5.**
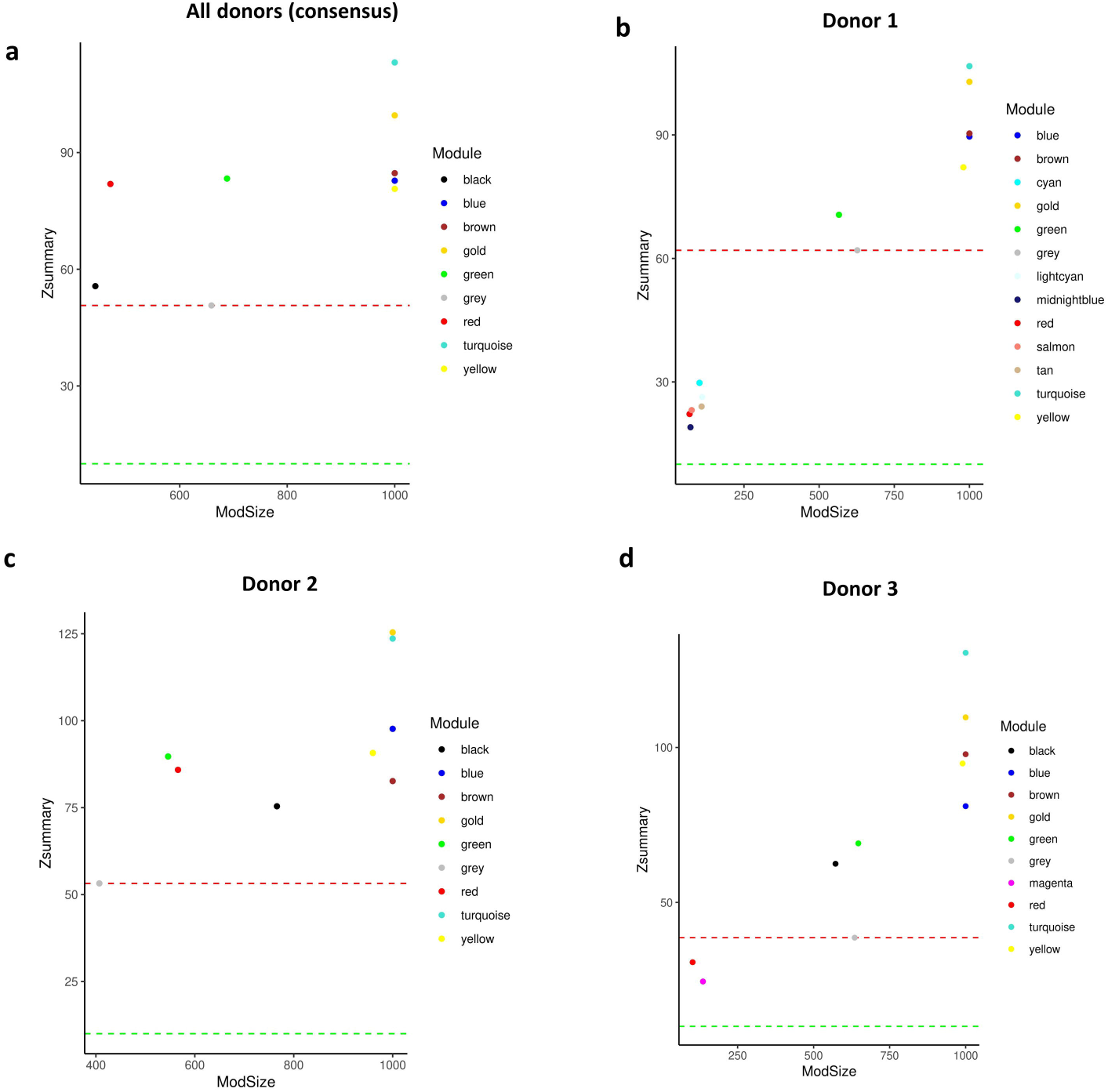
**a-d.** Zsummary plots for WGCNA modules for consensus modules **(a)** and individual donor modules **(b-d)**. High Zsummary values represent high stability. The red dotted lines represent the Zsummary intercept of the grey module (genes which failed to be classified). The green dotted line represents Zsummary=10, he minimal recommended threshold for evidence of conservation.

**Extended Data Figure 6.**
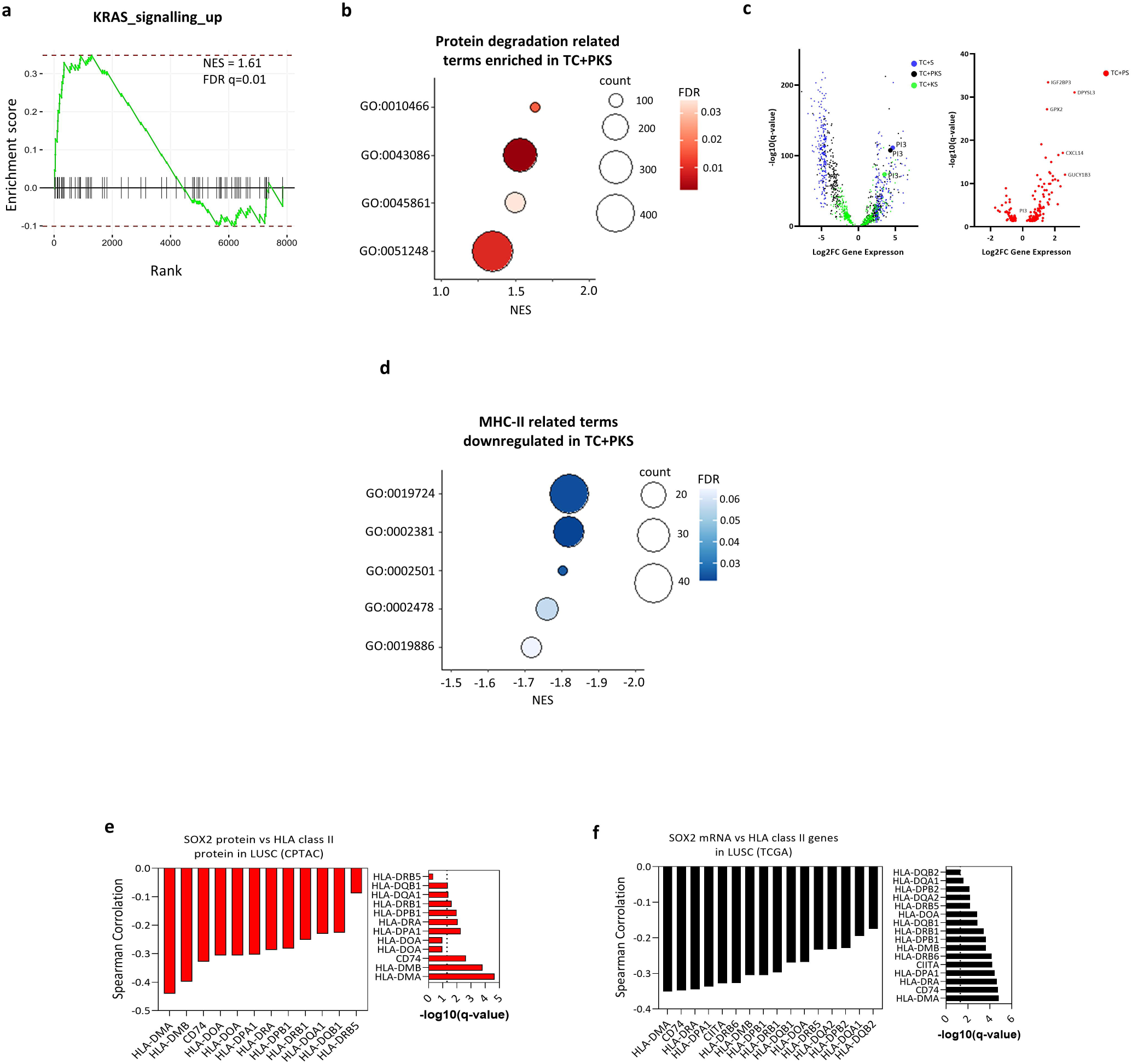
**a.** Gene set enrichment analysis showing the significant positive enrichment (q-value<0.05) of the KRAS signalling up hallmark (MSigDB, H: hallmark gene sets) in genes differentially expressed in TC+PKS versus TC (q-value<0.05). **b.** Gene set enrichment analysis with dot plot showing significant (q-value<0.05) positively enriched gene ontology terms related to protein degradation within genes differentially expressed (q-value < 0.05, log2FC in the TC+PKS versus TC comparisons. GO:0010466=negative regulation of peptidase activity, GO:0043086=negative regulation of catalytic activity, GO:0045861=negative regulation of proteolysis, GO:0051248=negative regulation of protein metabolic process. **c.** Volcano plots of differentially expressed genes in all *SOX2*^OE^ mutants versus TC mutants showing *PI3* expression regulation. **d.** Gene set enrichment analysis with dot plot showing significant (q-value<0.05) negatively enriched gene ontology terms related to antigen processing and presentation within genes differentially expressed (q-value<0.05) in the TC+PKS versus TC comparisons. GO:0019886=antigen processing and presentation of exogenous peptide antigen via MHC class II, GO:0002501=peptide antigen assembly with MHC protein complex, GO:0002478=antigen processing and presentation of exogenous peptide antigen, GO:0002381=immunoglobulin production involved in immunoglobulin mediated immune response, GO:0019724=B cell mediated immunity. **e-f.** Spearman correlations of genes associated with MHC-II antigen progressing and presentation with SOX2. Analysis was performed in cBioportal using LUSC samples from the CPTAC (protein) (n=80) and TCGA (mRNA) (n= 511) datasets. Left panels show -log10 (q-value) and dashed line indicates significance thresholds (q-value<0.05). p-values were calculated using 2-sided t-tests and q-values were calculated using the Benjamini-Hochberg FDR correction procedure.

**Extended Data Figure 7.**
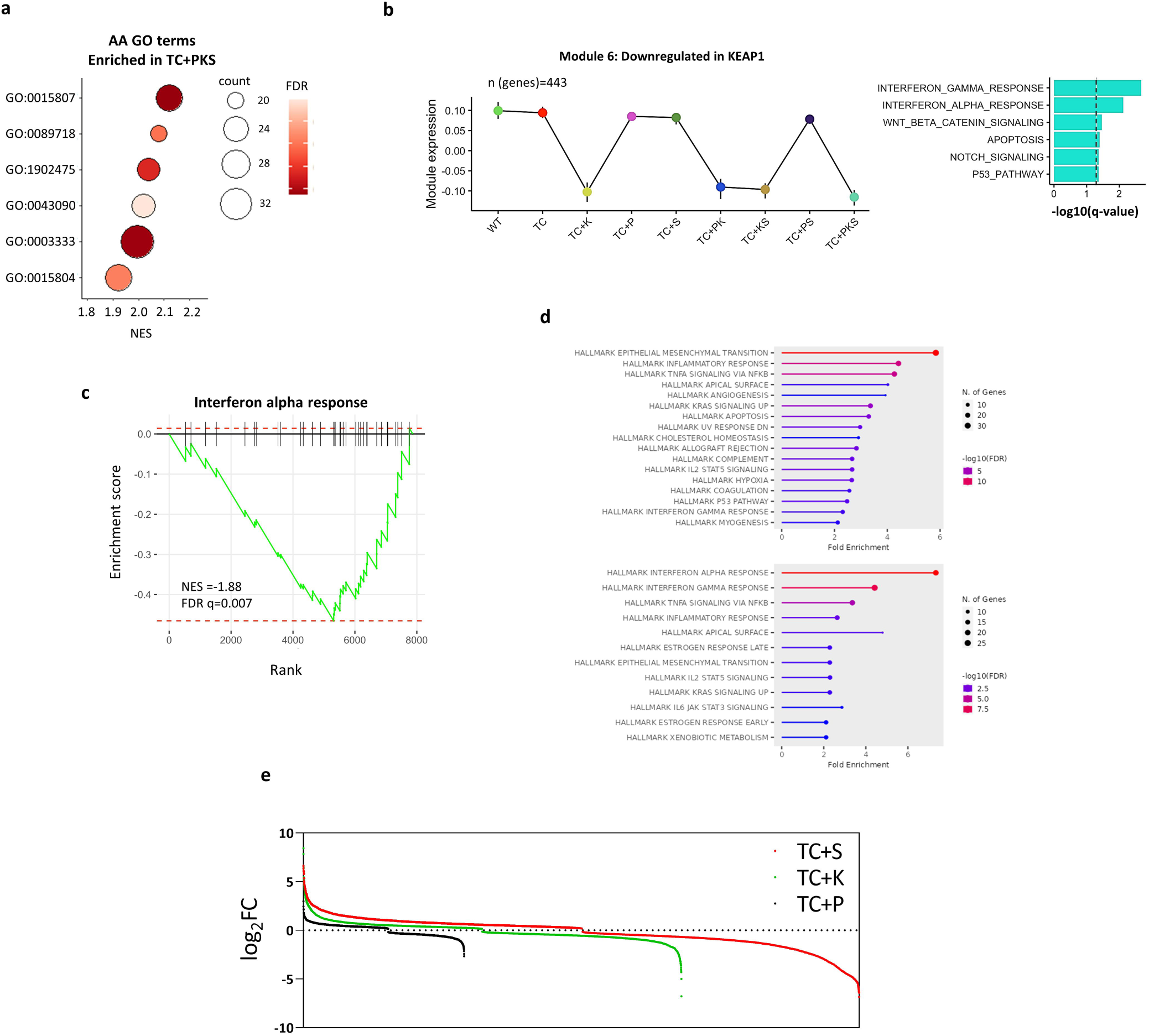
**a.** Gene set enrichment analysis with dot plot showing significant (q-value<0.05) positively enriched gene ontology terms related to amino acid transport within genes differentially expressed (q-value< 0.05) in the TC+PKS versus TC comparisons. GO:0015807=amino acid transport, GO:0003333=amino acid transmembrane transport, GO:1902475=alpha amino acid transmembrane transport, GO:0089718= amino acid import across plasma membrane, GO:0015804=neutral amino acid transport, GO:0043090= amino acid import. **b**. Left panel shows line graph depicting the expression of consensus module 6 in all hBEC mutants, showing that the expression dips in all KEAP1 disrupted mutants. Right panel shows overrepresentation analysis for hallmarks (MSigDB, H: hallmark gene sets), carried out using module 6 genes. Dotted line indicates significance threshold (q-value<0.05). **c.** Gene set enrichment analysis showing the negative enrichment of interferon alpha response (top) and interferon gamma response (bottom) hallmarks (MSigDB, H: hallmark gene sets) in genes differentially expressed in TC+PKS versus TC (q-value<0.05). **d.** Hallmarks enrichment (MSigDB, H: hallmark gene sets) in genes differentially expressed in LUSC with alterations targeting the OSR pathway (q>0.05). These include mutations or copy-number changes in *NFE2L2*, *KEAP1* and *CUL3*. Differentially expressed genes were downloaded from the cBioportal CPTAC (protein, top) and TCGA (mRNA, bottom) databases and enrichment analysis was carried out using the shinyGO application (FDR<0.05). **e.** Log2FC distributions of genes differentially expressed compared to TC mutants following activation of each of the three pathways (q-value<0.05).

## MATERIALS AND METHODS

### Cell culture

Mouse embryonic 3T3-J2 fibroblasts (Kerafast, EF3003) were cultured in DMEM (Gibco^TM^, 11995065) with 10% bovine serum (Fisher Scientific, 16-107-078), 2mM GlutaMAX™ Supplement (Gibco^TM^, 35050061) and 5% Penicillin-Streptomycin (Gibco^TM^, 15070063). Lenti-X 293T cells (Takara, 632180) and primary human lung fibroblasts (Lonza Bioscience, ID: HLF-2, batch:20TL356516) were cultured using DMEM (GibcoTM, 11995065) with 10% foetal bovine serum (LabTech, 80837), 2mM GlutaMAX™ Supplement (GibcoTM, 35050061) and 5% Penicillin-Streptomycin (GibcoTM, 15070063). Normal human bronchial epithelial cells (Lonza, CC-2540S) were selected based on age, negative smoking status, and absence of airway disease (supplementary table #) and cultured as described previously (1). Briefly, sub-confluent mouse embryonic 3T3-J2 fibroblasts were treated for 2 hours with 4µg/ml Mitomycin C (Sigma-Aldrich, M4287), washed three time with PBS, trypsinised (Gibco^TM^, 12605036), and seeded at 20,000 cells/cm^2^. Media was changed to a 1:3 ratio of Ham’s F-12 Nutrient Mix (Gibco^TM^, 11765054) to DMEM (Gibco^TM^, 41966) with 10% foetal bovine serum (Gibco^TM^, 11563397), 5µM Y-27632 Dihydrochloride (Bio-Techne Sales Corp, 1254), 25ng/ml hydrocortisone (Sigma-Aldrich. H0888), 0.125 ng/ml EGF (Thermo Fisher Scientific, 10-605-HNAE50), 5μg/ml insulin (Sigma-Aldrich, I6634), and 0.1nM cholera toxin (Sigma-Aldrich, C8052) and human bronchial epithelial cells were seeded at a density of approximately 20,000 cells/cm^2^. Fresh 3T3-J2 fibroblasts were prepared for every epithelial cell passage. All cells were cultured in a humidified incubator at 37°C with 5% CO_2_ and media was refreshed every two days and regularly tested for mycoplasma infection.

### Patient Samples

FFPE samples were obtained from the Manchester Cancer Research Centre (MCRC) Biobank. The MCRC Biobank holds a generic ethics approval (Ref: 18/NW/0092) which can confer this approval to users of banked samples via the MCRC Biobank Access Policy. Samples in this study were obtained through application 23_CALO_01.

### Plasmid construction

The one-step multiplex CRISPR-Cas9 assembly system kit was a gift from Takashi Yamamoto (Addgene Kit #1000000055) (2) was used to construct multiplex CRISPR-Cas9 plasmids used for the generation of cells harbouring knockouts in *TP53, CDKN2A*, *PTEN* and *KEAP1* genes. gRNAs were assembled into gRNA expression cassettes within pX330 plasmids using golden gate cloning, as described previously (2). For a comprehensive list of gRNAs and final pX330 plasmids see Supplementary Table 1. *SOX2* overexpression was achieved using the third-generation lentivirus vector, pUltra-hot, which was a gift from Malcom Moore (Addgene #24130). *SOX2* cDNA was amplified using restriction site PCR with 5’GAATTC (EcoRI) and 3’GAAGAC (BbsI) flanking sequences and ligated into the multiple cloning site of pUltrahot using a standard restriction digest and ligation, generating the pUltrahot-SOX2 vector.

### Bronchial epithelial cell electroporation and selection

Electroporation of multiplex CRISPR-Cas9 pX330 plasmids into normal bronchial epithelial cells was achieved using the P3 Primary Cell 4D-Nucleofector® X Kit L (Lonza, V4XP-3024) and 4D-Nucleofector™ X Unit (Lonza). 500,000 cells were pelleted and resuspended in 100µl Nucleofector™ Solution. 5µg of pX330 plasmid DNA was added and the full volume transferred to a Nucleocuvette™ vessel. Vessels were placed in the 4D-Nucleofector™ X Unit and electroporated using the DC-100 programme, incubated at room temperature for 10 minutes, and diluted with 500µl of pre-warmed media before seeding into a T25 containing 4.5ml of media and an adhered layer of mitotically inactivated mouse embryonic 3T3 fibroblasts, prepared as described previously (1). Cells were allowed to recover for two days, after which *TP53* mutant selection was achieved by addition of Nutlin-3A (Sigma-Aldrich, SML0580) to a final concentration of 5nM. Selection was carried out for 5 days, with media changes + Nutlin-3A every 2 days. Mutant bronchial epithelial cell colonies typically appeared 2-3 days following the end of selection.

### Bronchial epithelial cell transduction

The generation of mutant bronchial epithelial cells with stable *SOX2* overexpression was carried out by transduction using lentivirus particles harbouring the packaged pUltrahot-SOX2 vector. Lentivirus was generated by transfecting 90% confluent Lenti-X 293T cells with 880ng pUltrahot-SOX2, 572ng pMDLg/pRRE (Addgene #122251), 308ng pCMV-VSV-G (Addgene #8454), and 220ng pRSV-Rev (Addgene #12253) in 2ml of media and 6µl of FuGENE HD Transfection Reagent (Promega, #E2311). Virus titres were concentrated using a solution of polyethylene glycol (40% w/v) with NaCl (2.4% w/v) in sterile H_2_O. The number of virus copies/ml was determined using the Lenti-X™ qRT-PCR Titration Kit (Takara Bioscience, 631235) according to the manual. The same procedure was used to generate lentivirus particles harbouring the packaged pUltrahot vector lacking *SOX2* cDNA which served as an empty vector control. For transductions, 250,000 bronchial epithelial cells were seeded into a T25 containing 5ml of media and an adhered layer of mitotically inactivated mouse embryonic 3T3 fibroblasts, prepared as described previously (1). With bronchial epithelial cells still in suspension, a transduction mixture consisting of 4×10^8^ virus copies, 12mg/ml polybrene and PBS to 100µl was added to give a transduction efficiency of >90%. Transductions were carried out for 48 hours before fresh media was added, and initial transduction efficiencies were assessed by flow cytometry for mCherry detection.

### PCR genotyping and next generation amplicon sequencing

For genotyping to detect the presence of CRISPR-Cas9 mediated indels, DNA was extracted from mutant bronchial epithelial cells using proteinase K digestion. Approximately 500,000 cells were digested in 500µl of proteinase K digestion buffer (50mM KCl, 50 mM Tris-HCl, 2.5mM EDTA, 0.45% NP40 and 0.45% Tween) with 50µg Proteinase K (ThermoFisher Scientific, EO0491). Primers for PCR amplification were as follows: CDKN2A_forward 5’-CACCCTGGCTCTGACCATTC-3’, CDKN2A_reverse 5’-GCAAGTCCATTTCGGGATTA-3’, KEAP1_forward 5’-TACGACTGCGAACAGCGAC-3’, KEAP1_reverse 5’- GGCACAGAATCAAAGGTCAC-3’, PTEN_forward 5’- TCCAGTGTTTCTTTTAAATACCTGTT-3’, PTEN_reverse 5’- GGGGGAGAATAATAATTATGTGAGGT-3’, TP53_forward 5’- CAGGAAGCCAAAGGGTGAA-3’, and TP53_reverse 5’- CCCATCTACAGTCCCCCTTG-3’. PCR products were sequenced by Sanger sequencing and sequencing traces assessed for the presence of indels. For the assessment of % mutant reads by next generation amplicon sequencing, DNA was extracted using the DNeasy Blood & Tissue Kit (Qiagen, 69504) and target loci amplicons were generating by PCR using the above primers. Amplicons were then pooled to give a single amplicon admixture per mutant cell population. Targeted sequencing was carried out using the MiSeq Nano v2 Kit (Illumina) on the MiSeq desktop sequencer (illumina). Reads were aligned to hg38.

### Western blotting

Whole cell protein lysates were obtained by lysing cell pellets on ice in NP-40 buffer (150mM NaCl, 1% NP-40, 50mM tris pH 8) with 10x Protease Inhibitor Cocktail (Sigma-Aldrich, P8340). Protein quantification was done using the Pierce™ BCA Protein Assay Kit (Thermo Scientific, 23225). For sample preparation, 20-40ug of protein was mixed with 4x NuPAGE LDS buffer (Invitrogen, NP0007) and 10x NuPAGE reducing agent (Invitrogen, NP0004) and denatured at 90°C for 10 minutes. Equivalent amounts of each sample were run on 4-12% Bis-Tris gels (ThermoFisher Scientific, NP0322), transferred to methanol activated PVDF membranes and blocked for 1 hour at room temperature using 5% Marvel milk, or 5% BSA in TBST buffer. Membrane were immunoblotted with antibodies raised against p53 (SCB, sc-126) 1:1000, CDKN2A (p16) (SCB, 92803) 1:500, KEAP1 (SCB, sc-365626) 1:500, PTEN (SCB, sc-7974) 1:500, AKT (CST, 4691) 1:1000, pAKT(ser473) (CST, 4060) 1:1000, NQO1 (SCB, 32793) 1:1000, SOX2 (Abcam, ab97959) 1:1000, mCherry (CST, 43590) 1:1000, and Vinculin (Sigma, V9264) 1:10,000. Secondary antibodies were goat anti-rabbit IgG HRP (Agilent Technologies, P0440801-2), or rabbit anti-mouse IgG HRP (Agilent technologies, P044701-2) at 1:5000. Blots were developed using SuperSignal™ West Pico PLUS Chemiluminescent Substrate (ThermoFisher Scientific, 34577).

### Proliferation assays

Crystal violet cell proliferation assays were carried out in 6-well tissue culture plates. 50,000 bronchial epithelial cells were seeded into one well of a 12-well tissue culture plate containing an adhered layer of mitotically inactivated mouse embryonic 3T3 fibroblasts, prepared as described previously (1) and grown in a humidified incubator at 37°C with 5% CO2. Plates were harvest, fixed and stained at 24-hour intervals for 3 days. Briefly, 3T3-J2 cells were first removed with a quick trypsinisation, leaving bronchial epithelial cell colonies attached. Colonies were washed twice with ice-cold PSB and fixed for 10 minutes with ice-cold methanol. Colonies were then stained with 0.1% crystal violet dissolved in 20% methanol (w/w) in H_2_O for 15 minutes, de-stained under running water and dried overnight. Plates were imaged using an Epsom flatbed scanner and images analysed using Fiji (ImageJ) to give an output of percent confluency per well.

### Soft-agar colony forming assays

Colony forming assays were used to investigate anchorage independent growth. Base agar was generated by mixing a 1:1 ratio of 1% agar (w/v) in sterile water with cell culture media consisting of equal parts 2x DMEM (Gibco, 12800017) and 2x DMEM/F12 (Gibco, 12400024) supplemented with 20% FBS, 1uM Y-27632 (Bio-Techne, 1254/10), 0.5ng/ml EGF (Gibco, PHG0311), 1ug/ml hydrocortisone (Sigma-Aldrich, H0888-1G), 10ug/ml insulin (SLS, I6634), 0.2nM cholera toxin (Sigma-Aldrich, C8052), and 10% penicilin-steptomycin (Gibco, 15140122). Top agar was generated by mixing a 1:1 ratio of 0.7% agar (w/v) with cell culture media consisting of equal parts 2x DMEM (Gibco, 12800017) and 2x DMEM/F12 (Gibco, 12400024) with 10% sodium bicarbonate solution (v/v) (Gibco, 25080094). 1ml of base agar was added each well of a 6-well tissue culture plate and set for 5 minutes. A single cell suspension of 50,000 epithelial cells was mixed with 1ml top agar and seeded on top of the set base agar. Plates were incubated in a tissue culture incubator at 37°C with 5% CO_2_ for 4 weeks and fed with 300µl complete epithelial cell culture media once weekly. Colonies were visualised by staining with 0.005% crystal violet solution for 2-hours and 20x brightfield images of each well were captured using an inverted microscope. Colony quantification was carried out using Fiji where crystal violet-stained foci with a circumference >50µM were counted as a colony.

### Caspase-3/7 activity assays

Caspase-3/7 activity assays were carried out to determine the apoptotic effect of viral transduction using the pUltrahot-*SOX2* vector on mutant human bronchial epithelial cells. To capture apoptosis immediately after virus transduction, 3,200 bronchial epithelial cells were seeded into one well of a 96-well tissue culture plate containing an adhered layer of mitotically inactivated mouse embryonic 3T3 fibroblasts, prepared as described previously (1) with 50µl of media. With epithelial cells still in suspension, a transduction mixture consisting of 4×10^6^ virus copies, 12mg/ml polybrene, 5µM NucView 488 caspase-3 substrate (Biotium, 30029) and media to 50µl was added. Reversine (Sigma-Aldrich, R3904) was added to a final concentration of 2nM in positive control wells to induce caspase activity. Wells with transduction mixture containing equal amounts of the pUltrahot empty vector were used to control for background levels of virus mediated apoptosis. Plates were incubated in the Incucyte Live-Cell imager for 48 hours, with images captured every 60 minutes.

### Collagen-I invasion assays

Invasion assays were performed using human bronchial epithelial cells grown on collagen discs containing primary human pulmonary fibroblasts (Lonza Bioscience, ID: HLF-2, batch:20TL356516). Disc generation and contraction was carried out as described previously (3). For 12 discs, 1×10^6^ pulmonary fibroblasts were resuspended in 3ml FBS (Fisher Scientific, 16-107-078) and the whole volume added to a mixture of 20ml collagen-I (5mg/ml) (Enzo Life Sciences, ALX-522-435-0100), 5ml sterile H_2_O, 3ml 10x MEM (Gibco, 11430030), and 0.22M NaOH. 0.22M NaOH was added to increase the pH of the collagen mixture to achieve a transition from yellow to salmon pink (approximately pH 7). All components were prepared in a sterile cell culture hood and always kept on ice. 2.5ml of fibroblast-collagen mixture was added per 3mm^3^ cell culture dish and allowed to polymerise for 20 minutes in a 37°C humidified cell culture incubator. Following this, 1ml of prewarmed DMEM media (Gibco, 41966029) supplemented with 10% FBS, 5% penicillin-streptomycin (Gibco, 15070063) and 5% GlutaMAX (Gibco, 35050061) was added. The next day, an additional 1ml of media was added to each dish and media was changed every 2 days until the discs contracted to the size of one well of a 24-well tissue culture plate (approximately 8 days). Following contraction, collagen discs were transferred to a 24-well dish and 40,000 epithelial cells were seeded onto each disc in 1ml of PneumaCult™-Ex Plus expansion media (STEMCELL Technologies, 05040). Media was changed every 2 days for 1 week, after which each disc was suspended on a PET membrane in one well of a 6-well plate using tissue culture insert (Sarstedt, 83.3930). 2ml of complete PneumaCult™-ALI differentiation media (STEMCELL Technologies, 05001) was then added to the basal compartment of each well to generate an air-liquid interface, and media was changed every 2 days. Collagen-I invasion assays were cultured at 37°C in a humidified incubator with 5% CO_2_ for 3 weeks after which they were fixed overnight at 4°C in 4% paraformaldehyde, transferred to 70% ethanol and processed into FFPE blocks.

### Air-liquid interface culture and processing

Air liquid interface organotypic cultures were generated using human bronchial epithelial cells. Cultures were grown on Falcon® Permeable Supports with PET membranes (pore size 0.4µm) (Corning, 353095). Fistly, PET membranes were coated with 20µg type-I bovine collagen. Briefly, 3mg/ml PureCol® solution (CellSystems, 5005) was diluted 1:30 in sterile PBS and 200µl added to each PET membrane and incubated for 1 hour at room temperature. Collagen solution was removed, and membranes were washed gently with PBS. 40,000 bronchial epithelial cells were pelleted and resuspended in 100µl Airway Epithelial Cell Growth Medium (PromoCell, C-21160), after which the whole amount was seeded onto a collagen coated Falcon® Permeable Support placed within one well of a 24-wel plate. and 500µl of PneumoCult^TM^-ALI complete Basal Medium (STEMCELL Technologies, 05100) was added to the basal compartment. Epithelial cells were expanded for 7 days under submerged conditions with media changes every 2 days. On day 7, apical media was removed, and basal media replaced with 500µl of PneumoCult^TM^-ALI Complete Maintenance Medium (STEMCELL Technologies, 05100). Cultures were allowed to differentiate for 21 days with basal compartment media changes every 2 days. Once complete, cultures were process for downstream analysis. For histological assessment, cultures were fixed for 30 minutes in 4% paraformaldehyde, embedded in agarose, and processed into paraffin blocks for sectioning. For gene expression analysis, cultures were washed in PBS and RNA was extracted using the RNeasy Mini Kit (Qiagen, 74104) according to the manufacturer’s instructions.

### Immunofluorescence and immunohistochemistry

4µm FFPE sections were cut from FFPE blocks from ALI organotypic cultures, collagen-I invasion assays and clinical patient samples. Immunofluorescence staining was used to investigate mucociliary differentiation in ALIs by dual staining with anti-MUC5AC (Invitrogen, MA5-12178) (1:400) and anti-acetylated tubulin (Sigma, T6793) (1:400) antibodies and for identifying epithelial cells in invasion assays by dual staining for anti-vimentin (Invitrogen, MA5-11883) (1:250) and anti-EPCAM (Abcam, ab223582) (1/500). Antigen retrieval was performed in a pressurised vessel for 20 minutes at 95°C using Target Retrieval Solution, pH 9 (Agilent, S2367). 400µl/slide of primary antibody was incubated on slides overnight at 4°C. Secondary antibodies, including goat anti-mouse IgG1 Alexa Fluor 488 (Invitrogen, A-21121), goat anti-rabbit IgG Alexa Fluor 647 (Invitrogen, A32733), and goat anti-mouse IgG2b Alexa Fluor 647 (Invitrogen, A-21242), were diluted to 1:400 in PBS with 5% BSA and incubated on slides for 1 hour. Immunohistochemistry was used to investigate the expression of markers typically used to discriminate LUSC tumours from other lung cancers, including anti-TTF1 (Abcam, ab76013) (1:100), anti-cytokeratin 5/6 (Invitrogen, MA191106) (1:100) and anti-p63 (Abcam, ab124762) (1:400). The presence of club cells was investigated by immunohistochemistry using anti-CC10 (SCB, 365992) (1:2000), the expression of the squamous differentiation marker involucrin was done uing anti-involucrin (Invitrogen, MA5-11803) (1:200), SOX2 protein expression was investigated using anti-SOX2 (Abcam, ab92494) (1:100), and mCherry using anti-mCherry (Novus, NBP2-25157) (1:500). Antigen retrieval (ER 1, 20 minutes) and staining was performed using the Leica Bond RX automated platform using the classical IHC-F protocol with antigen retrieval achieved using BOND™ Epitope Retrieval solution 1 (Lecia, AR9961) for 20 minutes. CC10 and involucrin staining protocols included an additional 30-minute blocking step with casein (Sigma-Aldrich, B6429) and 5% bovine serum, respectively. Chromogenic 3,3-diaminobenzidine (DAB) staining was achieved using the BOND™ Polymer Refine Detection Kit (Lecia, DS9800).

### Image acquisition and analysis

All slides were scanned at 20x using the Olympus VS120 (fluorescence), or Olympus VS200 (brightfield). The images were analysed using HALO image analysis software (Indica Labs), with the Multiplex IHC 2.0 module. Percent positive MUC5AC, CC10, and p63 cells were calculated by dividing the number of positive cells by the total number of nuclei per analysis region. Percent acetylated tubulin coverage was calculated by dividing the length of acetylated tubulin positive epithelium by the total length of epithelium analysed. Fraction of mCherry positive epitheliums was calculated manually by dividing the length of mCherry positive epithelium by the total length of the epithelium. For invasion assays, measurements and manual counting of single invading epithelial cells (EPCAM^+^VIM^-^, or EPCAM^+^VIM^+^ cells), was achieved using the OlyVIA slide viewing software (Olympus). To calculate the number of invading cells per mm^2^, the number of invading cells was divided by the length of collagen disc cross-section. To avoid counting cells still attached to the main epithelial surface, EPCAM^+^VIM^-^, or EPCAM^+^VIM^+^ cells within 100µm of the epithelium base were discounted.

### Gene expression analysis by qPCR

qPCR was carried out using the Power SYBR Green PCR chemistry (Thermo Fisher Scientific, 4367659). cDNA was from total RNA using Superscript III (Thermo Fisher Scientific, 18080093) according to manufacturer’s instructions. The following oligonucleotides were used: SPRR2A (forward: gtatccaccgaagagcaagtaa, reverse: ggaacgaggtgagccaaata) and SPRR3 (forward: agcagaagaccaagcagaag, gacacagaaaacagatgggaaga) and beta-actin (forward: tggatcagcaagcaggagtatg, reverse: gcatttgcggtggacgat).

### RNA-sequencing

Air liquid interface organotypic cultures were washed once with PBS, snap frozen on dry ice and stored at -80°C for a maximum of 2-3 days. RNA was extracted using the RNeasy Mini kit (Qiagen, 74104) according to the manufacturer’s instructions. Specifically, each individual ALI culture was lysed using 200µl of RLT buffer. DNase digestion was performed on-column using the On-Column DNase I Digestion Set (Sigma Aldrich, DNASE70) as instructed. Finally, RNA was eluted in 30µl of RNase-free water, snap frozen on dry ice and stored at -80°C. Library preparation and RNA sequencing was carried out by the CRUK Manchester Institute Molecular Biology Core Facility. RNA quality was assessed using the Agilent 2100 Bioanalyzer and samples with a RIN value of >7 were selected for sequencing. Library preparation was performed using 100ng of total RNA using the Lexogen QuantSeq 3’ mRNA-Seq Library Prep Kit. Single-end sequencing with 100bp read length was performed using a Novaseq 6000 sequencer (Illumina). Basecalls were converted to fastq files using bcl2fastq (Illumina). Fastq files were trimmed for adapter sequences using the autodetect feature in trim_galore (version 0.6.5) and aligned to GRCh38 using Star aligner (version 2.5.1b). BAM alignments were quantified in R (version 3.6.1) using featureCounts from the Rsubread library (version 2.0.1). PCA plots were produced using regularised log-transformed read counts from DESeq2 (version 1.26). From CLG3, samples 07 and 11 were identified as outliers and removed from the analysis [1 from wild type and 1 from TC+P]. Due to the sequencing set-up, it was not possible to perform OLS regression for batch effect removal on the collected data. Surrogate variable analysis (SVA) was employed to identify and account for confounding covariates within the combined dataset. DESeq2 was used to perform comparisons between conditions within the combined dataset, including the SVA covariates in the model.

### Weighted gene co-expression network analysis

Data used for WGCNA were pre-processed from combined raw counts by filtering out low count genes (>10 reads allocated in 90% of samples). Raw counts were subsequently normalised using the vst function (DESeq2) (4) and surrogate variable effects removed using the removeBatchEffects function (limma) (5). To filter for only those genes differing between genotype, a mixed effects model (gene ∼ genotype), with donor info stated as a random effect, was compared to a null model (gene ∼ 1) using ANOVA; genes passing an adjusted p-value filter (pAdj < 0.05) were selected for further analysis. WGCNA (version 1.72-1) (6) was performed using dynamic power estimate optimisation (powers tested: 2-30). Gene-module allocations using the full dataset were used for further analyses (genes not allocated to a module reside within the “grey” module). Optimised power estimates were used for cross-validation with the sampledBlockwiseModules function (100 iterations of 80% samples across either 2000 or 80% of the input genes, depending on which was lower). Module stability/preservation statistics were calculated for the cross-validation results using the modulePreservation function. To assess heterogeneity between donors, WGCNA and cross-validation was also performed for each individual donor within each subset using a consistent power estimate for each donor. Gene expression within each module was plotted as heatmaps using scaled-normalised counts with the ComplexHeatmap package (version 1.72.1) (7). Cross-module comparisons were plotted using module eigengenes. Overenrichment analysis (ORA) of the module genes was performed using the clusterProfiler package (version 4.6.2) (8) with genesets from Hallmark signatures from MSigDB.

### Gene set enrichment analysis, overrepresentation analyses and publicly available LUSC sample databases

Preranked GSEA results were calculated for pairwise comparisons by fgsea (version 1.24.0) using the results defined by DESeq2, subsetting of genes by pAdj<0.05 and ranking using Log2FoldChange. Condensation of GO terms was performed using simplifyEnrichment (9).

Differentially expressed genes in LUSC patient samples with OSR-targeting alterations from the LUSC CPTAC (10) and TCGA (11) cohorts were downloaded from cBioportal (12) and enrichment analysis carried out using the shinyGO application (13).

Preinvasive lesion gene-expression data from Mascaux et al., (2019) (14) (GSE33479) was downloaded using the XTABLE application (15).

### Quantification and Statistics

Statistical testing of cell biology data including proliferation, colony forming, invasion assays, RT-qPCR, and the quantification of clinical sample and organoid immunostaining was carried out using the GraphPad Prism 9 software. Statistical comparisons were performed using a one-way ANOVA with *post hoc* tests for multiple comparison correction. In some cases, specific pairwise comparisons were selected from the larger dataset to increase power by reducing the number of comparisons made. These selections were justified in that they removed biologically meaningless, or redundant comparisons within the dataset. Significance was determined by p value < 0.05. Figures denote significance where ∗p < 0.05, ∗∗p < 0.01, ∗∗∗p < 0.001, and ∗∗∗∗p < 0.0001. Data was presented as means with standard deviation. n denotes the number of replicates

### Data and materials availability

RNAseq data has been uploaded to the GEO repository under the provisional project ID (PRJNA1043668).

## REFERENCES

1. Molina JR, Yang P, Cassivi SD, Schild SE, Adjei AA. Non-small cell lung cancer: epidemiology, risk factors, treatment, and survivorship. Mayo Clin Proc. 2008;83(5):584–94.

2. Chen Z, Fillmore CM, Hammerman PS, Kim CF, Wong KK. Non-small-cell lung cancers: a heterogeneous set of diseases. Nat Rev Cancer. 2014;14(8):535–46.

3. Ferone G, Song JY, Sutherland KD, Bhaskaran R, Monkhorst K, Lambooij JP, et al. SOX2 Is the Determining Oncogenic Switch in Promoting Lung Squamous Cell Carcinoma from Different Cells of Origin. Cancer Cell. 2016;30(4):519–32.

4. Wilkerson MD, Yin X, Hoadley KA, Liu Y, Hayward MC, Cabanski CR, et al. Lung squamous cell carcinoma mRNA expression subtypes are reproducible, clinically important, and correspond to normal cell types. Clin Cancer Res. 2010;16(19):4864–75.

5. Bray F, Ferlay J, Soerjomataram I, Siegel RL, Torre LA, Jemal A. Global cancer statistics 2018: GLOBOCAN estimates of incidence and mortality worldwide for 36 cancers in 185 countries. CA Cancer J Clin. 2018;68(6):394–424.

6. Paik PK, Pillai RN, Lathan CS, Velasco SA, Papadimitrakopoulou V. New Treatment Options in Advanced Squamous Cell Lung Cancer. Am Soc Clin Oncol Educ Book. 2019;39:e198–e206.

7. Siegel RL, Miller KD, Fuchs HE, Jemal A. Cancer statistics, 2022. CA Cancer J Clin. 2022;72(1):7–33.

8. Satpathy S, Krug K, Jean Beltran PM, Savage SR, Petralia F, Kumar-Sinha C, et al. A proteogenomic portrait of lung squamous cell carcinoma. Cell. 2021;184(16):4348–71 e40.

9. Cancer Genome Atlas Research N. Comprehensive genomic characterization of squamous cell lung cancers. Nature. 2012;489(7417):519–25.

10. Stewart PA, Welsh EA, Slebos RJC, Fang B, Izumi V, Chambers M, et al. Proteogenomic landscape of squamous cell lung cancer. Nat Commun. 2019;10(1):3578.

11. Hynds RE, Butler CR, Janes SM, Giangreco A. Expansion of Human Airway Basal Stem Cells and Their Differentiation as 3D Tracheospheres. Methods Mol Biol. 2019;1576:43–53.

12. Skoufou-Papoutsaki N, Adler S, D’Santos P, Mannion L, Mehmed S, Kemp R, et al. Efficient genetic editing of human intestinal organoids using ribonucleoprotein-based CRISPR. Dis Model Mech. 2023;16(10).

13. Cox JL, Wilder PJ, Desler M, Rizzino A. Elevating SOX2 levels deleteriously affects the growth of medulloblastoma and glioblastoma cells. PLoS One. 2012;7(8):e44087.

14. Wuebben EL, Wilder PJ, Cox JL, Grunkemeyer JA, Caffrey T, Hollingsworth MA, et al. SOX2 functions as a molecular rheostat to control the growth, tumorigenicity and drug responses of pancreatic ductal adenocarcinoma cells. Oncotarget. 2016;7(23):34890–906.

15. Timpson P, McGhee EJ, Erami Z, Nobis M, Quinn JA, Edward M, et al. Organotypic collagen I assay: a malleable platform to assess cell behaviour in a 3-dimensional context. J Vis Exp. 2011(56):e3089.

16. Behan FM, Iorio F, Picco G, Goncalves E, Beaver CM, Migliardi G, et al. Prioritization of cancer therapeutic targets using CRISPR-Cas9 screens. Nature. 2019;568(7753):511-6.

17. Tsherniak A, Vazquez F, Montgomery PG, Weir BA, Kryukov G, Cowley GS, et al. Defining a Cancer Dependency Map. Cell. 2017;170(3):564–76 e16.

18. Langfelder P, Horvath S. WGCNA: an R package for weighted correlation network analysis. BMC Bioinformatics. 2008;9:559.

19. Jiang Y, Jiang YY, Xie JJ, Mayakonda A, Hazawa M, Chen L, et al. Co-activation of super-enhancer-driven CCAT1 by TP63 and SOX2 promotes squamous cancer progression. Nat Commun. 2018;9(1):3619.

20. Wiedow O, Schroder JM, Gregory H, Young JA, Christophers E. Elafin: an elastase-specific inhibitor of human skin. Purification, characterization, and complete amino acid sequence. J Biol Chem. 1990;265(25):14791–5.

21. Cui C, Chakraborty K, Tang XA, Zhou G, Schoenfelt KQ, Becker KM, et al. Neutrophil elastase selectively kills cancer cells and attenuates tumorigenesis. Cell. 2021;184(12):3163–77 e21.

22. Mascaux C, Angelova M, Vasaturo A, Beane J, Hijazi K, Anthoine G, et al. Immune evasion before tumour invasion in early lung squamous carcinogenesis. Nature. 2019;571(7766):570-5.

23. Roberts M, Ogden J, Hossain ASM, Chaturvedi A, Kerr ARW, Dive C, et al. Interrogating the precancerous evolution of pathway dysfunction in lung squamous cell carcinoma using XTABLE. Elife. 2023;12.

24. Planells-Cases R, Lutter D, Guyader C, Gerhards NM, Ullrich F, Elger DA, et al. Subunit composition of VRAC channels determines substrate specificity and cellular resistance to Pt-based anti-cancer drugs. EMBO J. 2015;34(24):2993–3008.

25. Zavitsanou AM, Pillai R, Hao Y, Wu WL, Bartnicki E, Karakousi T, et al. KEAP1 mutation in lung adenocarcinoma promotes immune evasion and immunotherapy resistance. Cell Rep. 2023;42(11):113295.

26. Jamal-Hanjani M, Wilson GA, McGranahan N, Birkbak NJ, Watkins TBK, Veeriah S, et al. Tracking the Evolution of Non-Small-Cell Lung Cancer. N Engl J Med. 2017;376(22):2109–21.

27. Hodis E, Torlai Triglia E, Kwon JYH, Biancalani T, Zakka LR, Parkar S, et al. Stepwise-edited, human melanoma models reveal mutations’ effect on tumor and microenvironment. Science. 2022;376(6592):eabi8175.

28. Matano M, Date S, Shimokawa M, Takano A, Fujii M, Ohta Y, et al. Modeling colorectal cancer using CRISPR-Cas9-mediated engineering of human intestinal organoids. Nat Med. 2015;21(3):256–62.

29. Drost J, van Jaarsveld RH, Ponsioen B, Zimberlin C, van Boxtel R, Buijs A, et al. Sequential cancer mutations in cultured human intestinal stem cells. Nature. 2015;521(7550):43-7.

30. Ambrosone CB, Zirpoli GR, Hutson AD, McCann WE, McCann SE, Barlow WE, et al. Dietary Supplement Use During Chemotherapy and Survival Outcomes of Patients With Breast Cancer Enrolled in a Cooperative Group Clinical Trial (SWOG S0221). J Clin Oncol. 2020;38(8):804–14.

31. Omenn GS, Goodman GE, Thornquist MD, Balmes J, Cullen MR, Glass A, et al. Effects of a combination of beta carotene and vitamin A on lung cancer and cardiovascular disease. N Engl J Med. 1996;334(18):1150–5.

32. Beaulieu ME, Jauset T, Masso-Valles D, Martinez-Martin S, Rahl P, Maltais L, et al. Intrinsic cell-penetrating activity propels Omomyc from proof of concept to viable anti-MYC therapy. Sci Transl Med. 2019;11(484).

33. Arbour KC, Jordan E, Kim HR, Dienstag J, Yu HA, Sanchez-Vega F, et al. Effects of Co-occurring Genomic Alterations on Outcomes in Patients with KRAS-Mutant Non-Small Cell Lung Cancer. Clin Cancer Res. 2018;24(2):334–40.

34. Singh A, Daemen A, Nickles D, Jeon SM, Foreman O, Sudini K, et al. NRF2 Activation Promotes Aggressive Lung Cancer and Associates with Poor Clinical Outcomes. Clin Cancer Res. 2021;27(3):877–88.

35. van Boerdonk RA, Sutedja TG, Snijders PJ, Reinen E, Wilting SM, van de Wiel MA, et al. DNA copy number alterations in endobronchial squamous metaplastic lesions predict lung cancer. Am J Respir Crit Care Med. 2011;184(8):948–56.

36. McCaughan F, Pole JC, Bankier AT, Konfortov BA, Carroll B, Falzon M, et al. Progressive 3q amplification consistently targets SOX2 in preinvasive squamous lung cancer. Am J Respir Crit Care Med. 2010;182(1):83–91.

## Method References

1. Hynds RE, Butler CR, Janes SM, Giangreco A. Expansion of Human Airway Basal Stem Cells and Their Differentiation as 3D Tracheospheres. Methods Mol Biol. 2019;1576:43–53.

2. Sakuma T, Nishikawa A, Kume S, Chayama K, Yamamoto T. Multiplex genome engineering in human cells using all-in-one CRISPR/Cas9 vector system. Sci Rep. 2014;4:5400.

3. Timpson P, McGhee EJ, Erami Z, Nobis M, Quinn JA, Edward M, et al. Organotypic collagen I assay: a malleable platform to assess cell behaviour in a 3-dimensional context. J Vis Exp. 2011(56):e3089.

4. Love MI, Huber W, Anders S. Moderated estimation of fold change and dispersion for RNA-seq data with DESeq2. Genome Biol. 2014;15(12):550.

5. Ritchie ME, Phipson B, Wu D, Hu Y, Law CW, Shi W, et al. limma powers differential expression analyses for RNA-sequencing and microarray studies. Nucleic Acids Res. 2015;43(7):e47.

6. Langfelder P, Horvath S. WGCNA: an R package for weighted correlation network analysis. BMC Bioinformatics. 2008;9:559.

7. Gu Z, Eils R, Schlesner M. Complex heatmaps reveal patterns and correlations in multidimensional genomic data. Bioinformatics. 2016;32(18):2847–9.

8. Wu T, Hu E, Xu S, Chen M, Guo P, Dai Z, et al. clusterProfiler 4.0: A universal enrichment tool for interpreting omics data. Innovation (Camb). 2021;2(3):100141.

9. Gu Z, Hubschmann D. simplifyEnrichment: A Bioconductor Package for Clustering and Visualizing Functional Enrichment Results. Genomics Proteomics Bioinformatics. 2023;21(1):190–202.

10. Satpathy S, Krug K, Jean Beltran PM, Savage SR, Petralia F, Kumar-Sinha C, et al. A proteogenomic portrait of lung squamous cell carcinoma. Cell. 2021;184(16):4348–71 e40.

11. Cancer Genome Atlas Research N. Comprehensive genomic characterization of squamous cell lung cancers. Nature. 2012;489(7417):519–25.

12. Gao J, Aksoy BA, Dogrusoz U, Dresdner G, Gross B, Sumer SO, et al. Integrative analysis of complex cancer genomics and clinical profiles using the cBioPortal. Sci Signal. 2013;6(269):pl1.

13. Ge SX, Jung D, Yao R. ShinyGO: a graphical gene-set enrichment tool for animals and plants. Bioinformatics. 2020;36(8):2628–9.

14. Mascaux C, Angelova M, Vasaturo A, Beane J, Hijazi K, Anthoine G, et al. Immune evasion before tumour invasion in early lung squamous carcinogenesis. Nature. 2019;571(7766):570-5.

15. Roberts M, Ogden J, Hossain ASM, Chaturvedi A, Kerr ARW, Dive C, et al. Interrogating the precancerous evolution of pathway dysfunction in lung squamous cell carcinoma using XTABLE. Elife. 2023;12.

